# Estimating genetic correlation jointly using individual-level and summary-level GWAS data

**DOI:** 10.1101/2021.08.18.456908

**Authors:** Yiliang Zhang, Youshu Cheng, Yixuan Ye, Wei Jiang, Qiongshi Lu, Hongyu Zhao

## Abstract

With the increasing accessibility of individual-level data from genome wide association studies, it is now common for researchers to have individual-level data of some traits in one specific population. For some traits, we can only access public released summary-level data due to privacy and safety concerns. The current methods to estimate genetic correlation can only be applied when the input data type of the two traits of interest is either both individual-level or both summary-level. When researchers have access to individual-level data for one trait and summary-level data for the other, they have to transform the individual-level data to summary-level data first and then apply summary data-based methods to estimate the genetic correlation. This procedure is computationally and statistically inefficient and introduces information loss. We introduce GENJI (Genetic correlation EstimatioN Jointly using Individual-level and summary data), a method that can estimate within-population or transethnic genetic correlation based on individual-level data for one trait and summary-level data for another trait. Through extensive simulations and analyses of real data on within-population and transethnic genetic correlation estimation, we show that GENJI produces more reliable and efficient estimation than summary data-based methods. Besides, when individual-level data are available for both traits, GENJI can achieve comparable performance than individual-level data-based methods. Downstream applications of genetic correlation can benefit from more accurate estimates. In particular, we show that more accurate genetic correlation estimation facilitates the predictability of cross-population polygenic risk scores.

## Introduction

Genetic correlation analysis, which quantifies the correlation of additive genetic effects on different traits across a set of genetic markers, has gained popularity owing to the remarkable success of genome-wide association study (GWAS) in the past 15 years^1^. Compared with traditional family-based approaches^2^, estimating genetic correlation from GWAS data does not require samples from large pedigrees. The phenotypes of interest also do not have to be measured on the same individuals. These advances have effectively increased the sample size and improved statistical power in genetic correlation studies. Consequently, genetic correlation estimation has become a routine for post-GWAS analysis^3^ and has shed light on the shared genetic basis of numerous human complex traits and diseases^4,5^.

A plethora of GWAS-based methods for genetic correlation estimation have been developed^6^, such as genome-based restricted maximum likelihood^7^ (GREML) for individual-level data and linkage disequilibrium (LD) score regression^8^ (LDSC) for summary-level data. The GREML approach is based on variance components estimation in a linear mixed model (LMM) framework which needs individual-level genotype and phenotype data as input. Many computational tools have been developed to implement GREML^9–11^, which mainly differ by the algorithms used for likelihood optimization. LDSC, in comparison, only requires GWAS summary data and is thus more broadly used in GWAS analyses. Web servers have also been built to facilitate the computation and visualization of genetic correlations^12^.

Despite the popularity, estimates of LDSC are substantially less precise with larger standard errors compared to GREML^6,13^ due to information loss from individual-level data to summary statistics. This suggests that when individual-level data are available for both traits, GREML (or similar methods based on individual-level data) should be the method of choice. In addition, transethnic genetic correlation is recently introduced to quantify how the genetic architecture of complex traits varies across populations^14^. Summary statistics-based approaches are particularly less reliable for transethnic genetic correlation due to typically smaller sample sizes of non-European GWAS and a lack of robustness to mismatched LD between GWAS samples and reference panels. Even for analysis of GWAS from the same population, summary statistics-based methods struggle when the input GWAS have limited power.

Currently, if individual-level data are available for a trait of interest, in order to test its genetic correlations with published GWAS, researchers would have to first generate summary data for the trait and then estimate genetic correlations based on the summary statistics. Such a procedure is conspicuously inefficient. Our main goal in this study is to find a way to jointly model individual-level data (e.g., for a disease with limited samples or a GWAS in minority populations) and summary statistics (e.g., from published meta-analyses) to efficiently estimate genetic correlation.

We introduce GENJI (Genetic correlation EstimatioN Jointly using Individual-level and summary data), a method that estimates genetic correlation with individual-level data for one trait and summary-level data for the other trait. Through extensive simulations, we demonstrate that GENJI provides statistically rigorous and computationally efficient inference for both within-population and transethnic genetic correlations and substantially outperforms summary data-based methods. We demonstrate the effectiveness of GENJI through applications to many datasets, including UK Biobank (UKBB)^15^, the Wellcome Trust Case Control Consortium (WTCCC)^16^, the Northern Finland Birth Cohorts program (NFBC), and numerous GWAS of complex human traits spanning European, African, and East Asian populations from published studies and BioBank Japan (BBJ)^17,18^.

## Results

### Overview of GENJI

Genetic correlation (covariance) is the correlation (covariance) of additive genetic components of complex traits across a set of genetic markers. It is commonly used as an informative metric to quantify the shared genetic basis between two traits and has also been extended to study the shared and unique genetic effects on the same trait in multiple populations. In this paper, we refer to the former as within-population genetic correlation and the latter as transethnic genetic correlation. In what follows, without stated explicitly, within-population genetic correlation and transethnic genetic correlation are collectively referred to as genetic correlation. The proposed method can estimate both types of genetic correlations.

Genetic correlation *r_g_* is genetic covariance *ρ_g_* normalized by SNP heritability, i.e., 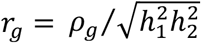, where 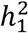 and 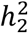 are the heritability of the two traits, respectively. Assume that we have individual-level GWAS data of the first trait (referred to as study 1 in the following) but only summary-level data of the second trait (referred to as study 2 in the following). We model phenotype *ϕ* as the sum of genetic component *Xβ* and random noise, where *X* and *β* represent genotype matrix and genetic effects, respectively. Denote *z*_2_ as the vector of z-scores in study 2. The LD matrix for study 2 can be estimated by an external reference panel or even genotype data in study 1 if they are conducted on the same population. We compute expectations and variances of 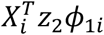 and apply weighted least squares to estimate genetic covariance:

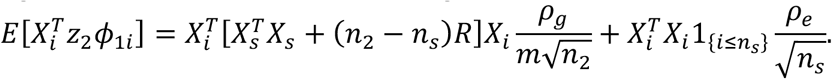

Instead of block jackknife used in LDSC^8^, the estimates of GENJI can use the theoretical standard error from weighted regression because individuals in study 1 are presumably unrelated to each other (**Methods**).

In practice, there can be some individuals shared by the two studies. Although we have individual-level data for the overlapped samples in study 1, we assume the phenotypes of them for study 2 are unknown. We assume the first *n_s_* samples in both studies are the overlapped samples. Similar to LDSC^8^, sample overlap inflates the covariance between *ϕ*_1*i*_ and 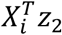 for each overlapped individual. As a result of the expectation of 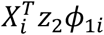, genetic covariance *ρ_g_* and noise covariance *ρ_e_* appear in different terms and hence can be distinguished by regression (**Methods**).

### Simulations for within-population genetic covariance

We performed simulations to assess the performance of GENJI on within-population genetic covariance estimation. We compared GENJI with two state-of-the-art methods, LDSC and GREML, on both quantitative and binary traits. We provided GREML with individual-level data, LDSC with summary-level data, and GENJI with individual-level data for the first study while summary-level data for the second one.

To assess the robustness of different methods to sample overlap, we changed the proportion of overlapping samples from zero to half of the sample sizes of the studies. We used real genotype data from WTCCC^16^ to simulate traits. We equally divided 15,918 samples in the dataset into two subsets which we denote as set 1 and set 2, respectively. There is no shared individuals between set 1 and set 2. Randomly combining the samples from set 1 and set 2, we created set 3, set 4, and set 5 which had 10%, 25%, and 50% overlapping samples with set 1, respectively. The simulated GWASs were conducted on these subsets and the sample sizes of them were all equal to 7,959. The SNP effects were generated by a bivariate normal distribution with genetic covariance ranging from 0 to 0.25 and heritability was fixed at 0.5. Each setting was repeated 100 times. Detailed simulation settings and quality control procedures are described in the **Methods** section.

The three competing methods in this section showed well-calibrated type I error rates when the true covariance was zero (**Supplementary Figures 1-4**). LDSC and GENJI provided unbiased estimates across all the settings. GREML overestimated genetic covariance for the 10% and 25% sample overlap settings (**Figure 1**).

**Figure 1.**
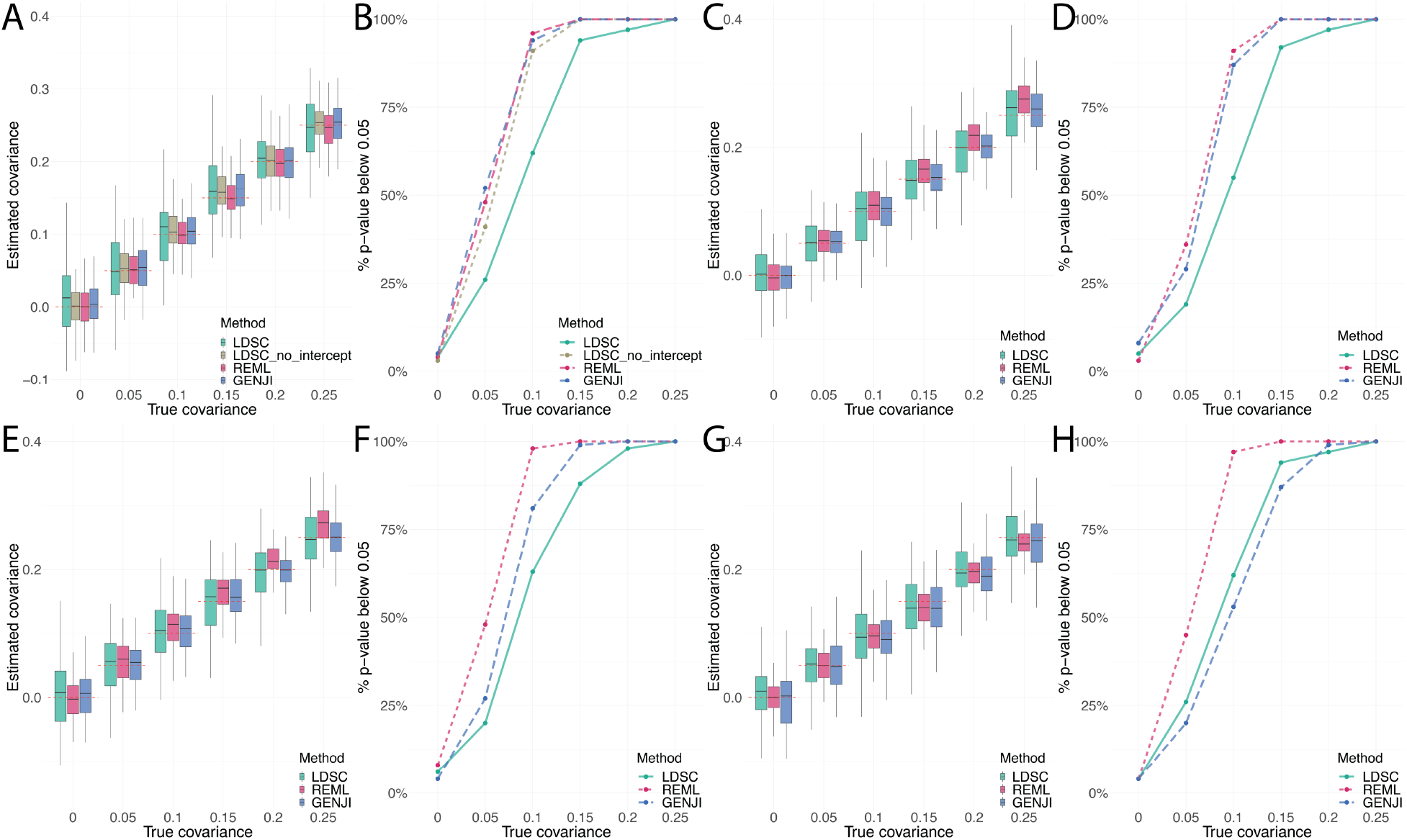
Simulation results for within-population genetic covariance estimation. We compare the performance of LDSC, GREML, and GENJI by point estimation, type-I error, and statistical power. Boxplots show the quantiles of the estimates of genetic covariance in different settings. The red dashed lines represent the true value of genetic covariance. We use the proportion of p-values that are less than 0.05 to estimate type-I error or statistical power when true parameters are zero or nonzero, respectively. **(A-B)** Two GWASs are simulated on two non-overlapping datasets (set 1 and set 2). Because there is no shared individual between the studies, we also include LDSC that constrain the intercept to zero in the comparison and denote it as “LDSC_no_intercept”. Panels C-H compare LDSC, GREML, and GENJI using GWASs **(C-D)** with a 10% sample overlap (set 1 and set 3), **(E-F)** with a 25% sample overlap (set 1 and set 4), and **(G-H)** with a 50% sample overlap (set 1 and set 5).

In the case of zero overlapping sample (set 1 and set 2), we also included LDSC with intercept fixed at zero (denoted as “LDSC_no_intercept”) in the comparison. When there was no overlapping sample, LDSC_no_intercept, GREML and GENJI had comparable estimates while the statistical power of GREML or GENJI was larger than that of LDSC_no_intercept when the true value of genetic covariance was relatively low. Estimates of LDSC without constraint had substantially larger variance and lower power than the other methods (**Figure 1A-B**). This indicates that although no extra information for point estimation of genetic covariance can be acquired from individual-level data compared with summary-level data for studies with disjoint cohorts, the standard errors from parametric methods are more stable than those from resampling-based methods (e.g., block jackknife). When we did not fix intercept for LDSC, the variability of intercept estimates increased the standard error of genetic covariance by 30% compared to LDSC with a constraint, a phenomenon that was also reported in the paper of LDSC^8^.

For simulations in set 1 and set 3 (10% overlap), set 4 (25% overlap), and set 5 (50% overlap), GENJI outperformed LDSC when the overlapping samples were less than half of the sample sizes of the studies (**Figure 1C-F**) and achieved comparable results with LDSC when the proportion of overlapping samples was 50% (**Figure 1G-H**). GENJI was even as powerful as GREML and had more accurate estimates under low sample overlap setting (<10% sample overlap). Under moderate sample overlap setting (25%), the power of GENJI was in between LDSC and GREML. We observe that the advantage of GENJI over LDSC reduces as the overlapped sample size increases, and reaches zero when half of study 1 samples are included in study 2.

We also performed simulations for binary traits and obtained similar results except that genetic covariance is on the observed scale (**Supplementary Figure 5**).

### Simulations for transethnic genetic correlation

We performed simulations to assess the performance of GENJI on transethnic genetic covariance estimation. We compared GENJI with Popcorn^14^ and GREML. In this simulation, the first GWAS was of African ancestry and the second was of European ancestry. Genotypes of 6,992 related samples of African ancestry from UKBB were used to simulate the phenotypes of study 1. We still used samples from WTCCC (set 1; n=7,959) to simulate phenotypes of European ancestry for study 2. The SNP effects were generated by a bivariate normal distribution with genetic covariance ranging from 0 to 0.5 and heritability was fixed at 0.5. We note that when genetic covariance is 0.5, the genetic correlation is equal to 1 which means the genetic effects are perfectly correlated between the populations. Each simulation setting was repeated 100 times. Detailed simulation settings and quality control procedures are described in the **Methods** section.

Popcorn needs two external reference panels to estimate LD matrices in two populations. To make fair comparisons, we also added an additional setting for Popcorn using in-sample African references, which are also the African genotype data used for simulations (denoted as “Popcorn_in_sample”). We included two kinds of hypothesis testing: the null hypothesis is either (1) genetic correlation = 0; or (2) genetic correlation = 1.

GENJI outperformed both Popcorn using external reference panel and Popcorn using in-sample reference panel, showing more accurate estimates and improved statistical power (**Figure 2**). Due to constraint of genetic correlation being less than 1 for GREML, GREML under-estimated genetic covariance for the setting of 0.5. GENJI showed unbiased estimates across all the settings (**Figure 2A**). Estimates of GENJI were as powerful as those of GREML in most settings (**Figure 2B-C**). All methods showed well-controlled type-I error for both kinds of testing (**Supplementary Figure 6-7**).

**Figure 2.**
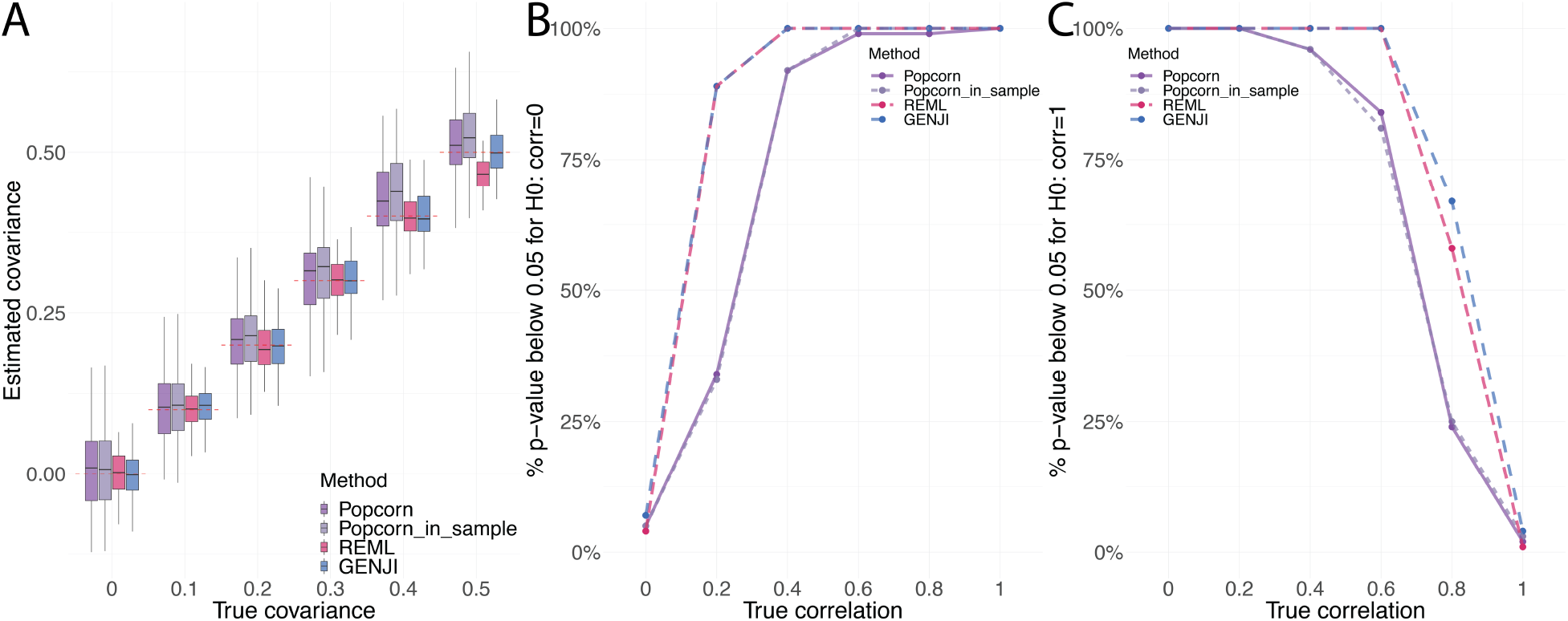
Simulation results for transethnic genetic covariance estimation. We compare the performance of Popcorn, GREML and GENJI by point estimation, type-I error and statistical power. We include Popcorn using both external and in-sample reference panel and denote the later as “Popcorn_in_sample”. **(A)** Boxplots show the quantiles of the estimates of genetic covariance in different settings. The red dashed lines represent true value of genetic covariance. **(B-C)** We use the proportion of p-values that are less than 0.05 to estimate type-I error or statistical power under true null or false null, respectively.

**Figure 3.**
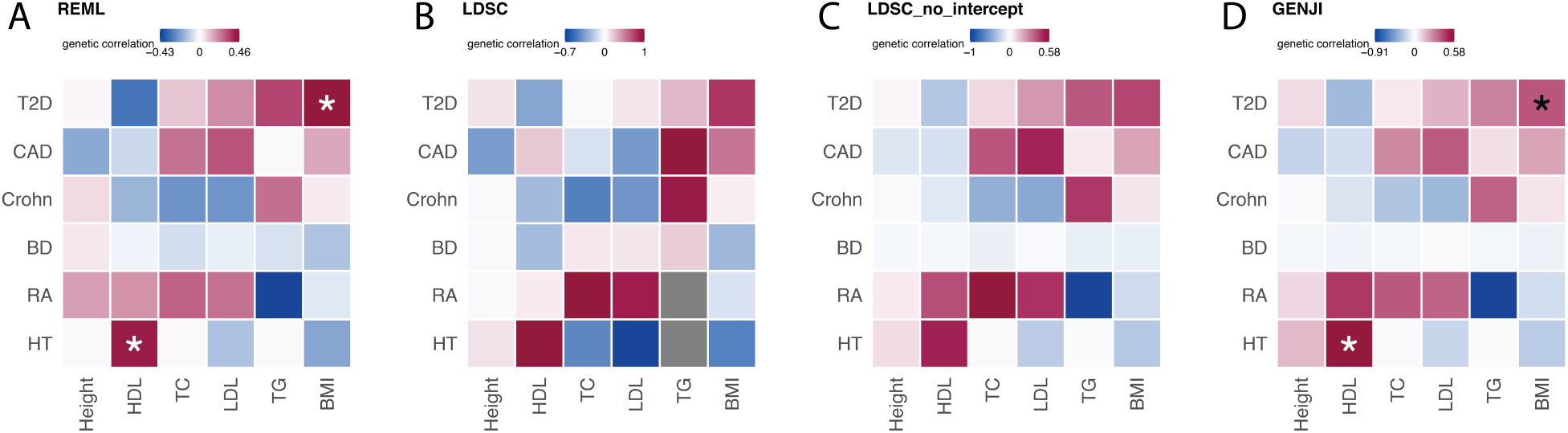
Genetic correlations estimated by GREML, LDSC, and GENJI across traits in WTCCC and NFBC. The heatmaps reflect the genetic correlation estimates of **(A)** GREML, **(B)** LDSC, **(C)** LDSC with intercept fixed at zero, and **(D)** GENJI. Asterisks in the heatmap highlight significant genetic correlations after Bonferroni correction for 36 pairs. Due to low heritability estimation, estimates of LDSC for genetic correlation of two trait pairs are unavailable and colored as grey in panel (B). We summarize detailed information about each trait, including abbreviations, in **Supplementary Tables 1** and **2**.

### Real data applications: within-population genetic correlation

We applied GREML, LDSC, and GENJI to estimate genetic correlations across 12 traits in WTCCC (n~16,000) and NFBC (n~5,300) (**Methods**). Both studies were conducted on samples of European ancestry and shared zero samples hence LDSC_no_intercept can also be applied. Detailed information about the GWAS data is summarized in **Supplementary Tables 1** and **2**. Since individual-level data of both studies are available, GREML can be treated as the gold standard for the other two methods. Consistent with our simulation results, genetic covariance and genetic correlation estimates of GENJI and LDSC_no_intercept were more consistent with GREML estimates, measured by R^2^, than those of LDSC without a constraint on intercept (**Supplementary Figure 8**). Although the point estimates of GENJI and LDSC_no_intercept on the 36 trait pairs were similar, GENJI achieved higher statistical power. After Bonferroni correction (p<0.05/36=1.39e-3), genetic correlations of two trait pairs were identified by both GREML and GENJI while LDSC or LDSC with intercept fixed at zero failed to identify any trait pairs (**Supplementary Table 3**). The significant trait pairs are type-2 diabetes (T2D)^19^ versus body mass index (BMI)^20^ and hypertension (HT) versus high-density lipoprotein (HDL)^21^. The positive association between obesity (BMI) and T2D has long been observed^22^ and is replicated in our following applications on GWAS summary data with larger sample sizes (**Figure 4**). Although low levels of HDL cholesterol are associated with increased risk of coronary artery disease (CAD)^23^, positive correlations between HDL and HT were observed across different populations^24,25^. In a recent study, the authors showed that the biological mechanism of the positive association between HDL and HT is related to circulating CD34-positive cell levels^26^.

To illustrate the superior power of GENJI over LDSC on more GWASs with larger sample sizes, we applied GENJI and LDSC to estimate the genetic correlations between 25 common traits with publicly available GWAS summary data and 6 traits from UKBB (n~270,000), which are the same set of the 6 NFBC traits we used before. We fixed the intercept of LDSC at zero because there are no overlapped samples between the trait pairs. We provided GENJI with the individual-level data from UKBB. Detailed information of the 25 GWAS summary data is summarized in **Supplementary Table 4**. Detailed information of the 6 traits from UKBB is summarized in **Supplementary Table 5**.

The genetic correlation estimates of GENJI and LDSC are highly consistent (R^2^=0.96). However, the standard errors from GENJI are mostly smaller than those from LDSC. Of the 150 trait pairs, significant genetic correlations of 82 and 33 trait pairs are identified by GENJI and LDSC, respectively, after Bonferroni correction (p<0.05/150=3.3e-4) (**Figure 4**; **Supplementary Table 6**). All the trait pairs identified by LDSC are also identified by GENJI.

**Figure 4.**
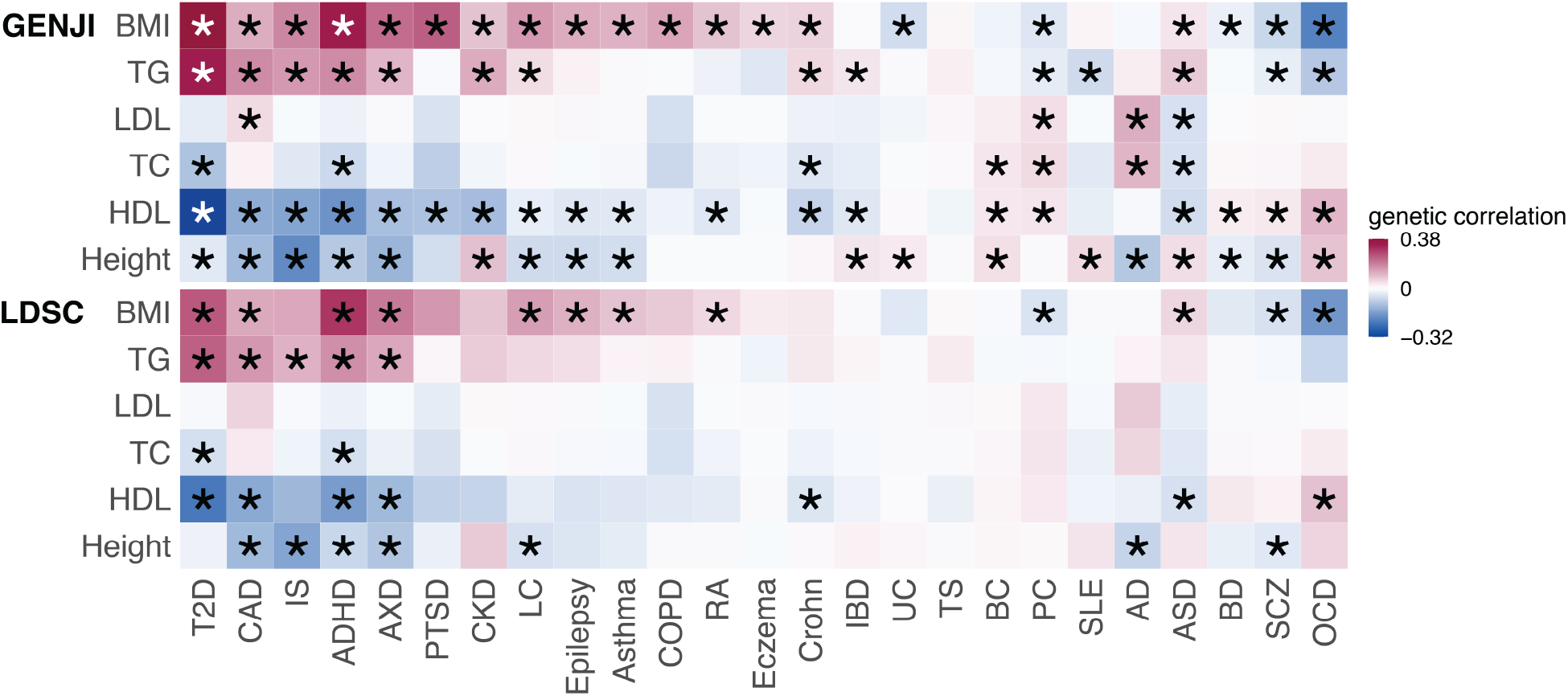
Genetic correlation estimates of GENJI and LDSC for 150 trait pairs. The upper and lower heatmaps reflect the estimates of GENJI and LDSC, respectively. Asterisks in the heatmap highlight significant genetic correlations after Bonferroni correction for the 150 pairs. We summarize detailed information about each trait, including abbreviations, in **Supplementary Tables 4** and **5**.

### Real data applications: transethnic genetic correlation

We applied GENJI and compared the performance of GENJI with Popcorn on transethnic genetic correlation estimation. We estimated the transethnic genetic correlation of 12 complex traits in the African, East Asian, and European populations. The choice of these traits was based on data availability and trait heritability (**Methods**). The GWAS data of African and European populations were from UKBB. The GWAS data of the East Asian population were from BBJ^17,18^ due to the small sample size of East Asian ancestry in UKBB (n~1,500). We only have access to summary-level GWAS data of BBJ. The details about the sample sizes and the sources of the data that we use in this section can be found in **Supplementary Tables 5** and **7**.

We first compared GENJI, Popcorn, and GREML on transethnic genetic correlation between European and African populations in UKBB. To reduce the computational burden for GREML, we used a subset of European ancestry samples from UKBB (n=10,000). For comparison fairness, GENJI and Popcorn also used the reduced dataset as input in this analysis. The individual-level data of African GWASs and summary-level data of European ancestry were provided to GENJI. We applied Popcorn twice in the comparison: (1) we used genotype data from the 1000 Genomes Project^27^ as the reference panels for African and European GWASs; (2) we used genotype data of African samples from UKBB as the reference panel for African GWASs (in-sample reference panel) and European samples from 1000 Genomes Project as the reference panel for European GWASs. It was also for comparison fairness because GENJI took advantage of the information from individual-level data of African ancestry. The results of the GREML, GENJI, and Popcorn are presented in **Supplementary Table 8**. The estimates of GENJI were substantially more consistent with GREML estimates (R^2^=0.94) compared to Popcorn (R^2^=0.41) or Popcorn_in_sample (R^2^=0.45) (**Supplementary Figure 9**). The variability of the Popcorn estimates yielded relatively unreliable and improper outcomes. To be specific, the transethnic genetic correlation estimates of Popcorn or Popcorn_in_sample for four traits were outside of [-1,1], which is the range of genetic correlation. Three and two traits were unavailable for Popcorn and Popcorn_in_sample, respectively, because of negative estimates for heritability. As a comparison, two traits’ GREML estimates were outside the proper range while no genetic correlation estimates of GENJI were out of bound. Estimates for all traits were available for GREML and GENJI. Consistent with our simulation results, standard errors of GENJI were substantially smaller than those of Popcorn and similar to those of GREML (**Supplementary Figure 10**). No traits were identified by Popcorn_in_sample with transethnic genetic correlation significantly larger than zero after Bonferroni correction (p<0.05/24=2.1e-3) while both GREML and GENJI identified three significant traits: Height, BMI and HDL.

We then used the whole dataset of European ancestry samples in UKBB and GWAS summary data from BBJ to estimate the transethnic genetic correlation across African, East Asian, and European populations for the 12 traits. The genotype data provided to GENJI were also used as the reference panel for Popcorn (in-sample reference panel) (**Methods**). The point estimates from GENJI and Popcorn were similar (**Figure 5**). Since the sample sizes of the GWAS summary data from BBJ were larger than those of African population GWASs (**Supplementary Table 5** and **7**), the standard errors for the transethnic genetic correlation between European and East Asian populations were smaller than the other two population pairs. The standard errors of GENJI estimates were consistently smaller than those of Popcorn. Due to the unstable estimates for heritability, the transethnic genetic correlation estimates of Popcorn for prostate cancer (PC)^28^ were unavailable. The genetic correlations between European and East Asian population were significantly larger those between European and African populations (Wilcoxon test p=4.9e-4) and those between East Asian and African populations (Wilcoxon test p=9.8e-4) across the 12 traits. The differences of genetic correlation between African and European populations and African and East Asian populations were not statistically significant (Wilcoxon test p=0.42). The transethnic genetic correlation estimates of GENJI for most traits were significantly larger than zero and smaller than one across the populations (**Supplementary Tables 9-11**), which indicates that the genetic effects for most traits are neither independent nor perfectly correlated between the populations. All traits show genetic correlation significantly larger than zero between European and East Asian populations according to the results of GENJI. Only two traits, age at natural menopause (ANM)^29^ and PC did not show genetic correlations significantly larger than zero between European and African populations. A recent transancestry GWAS of PC showed that the gap of R^2^ of the polygenic risk score (PRS) for African ancestry was substantially lower than that of East Asian and European ancestry if the polygenic risk score (PRS) is trained by the GWAS which mostly consists of European ancestry^30^.

**Figure 5.**
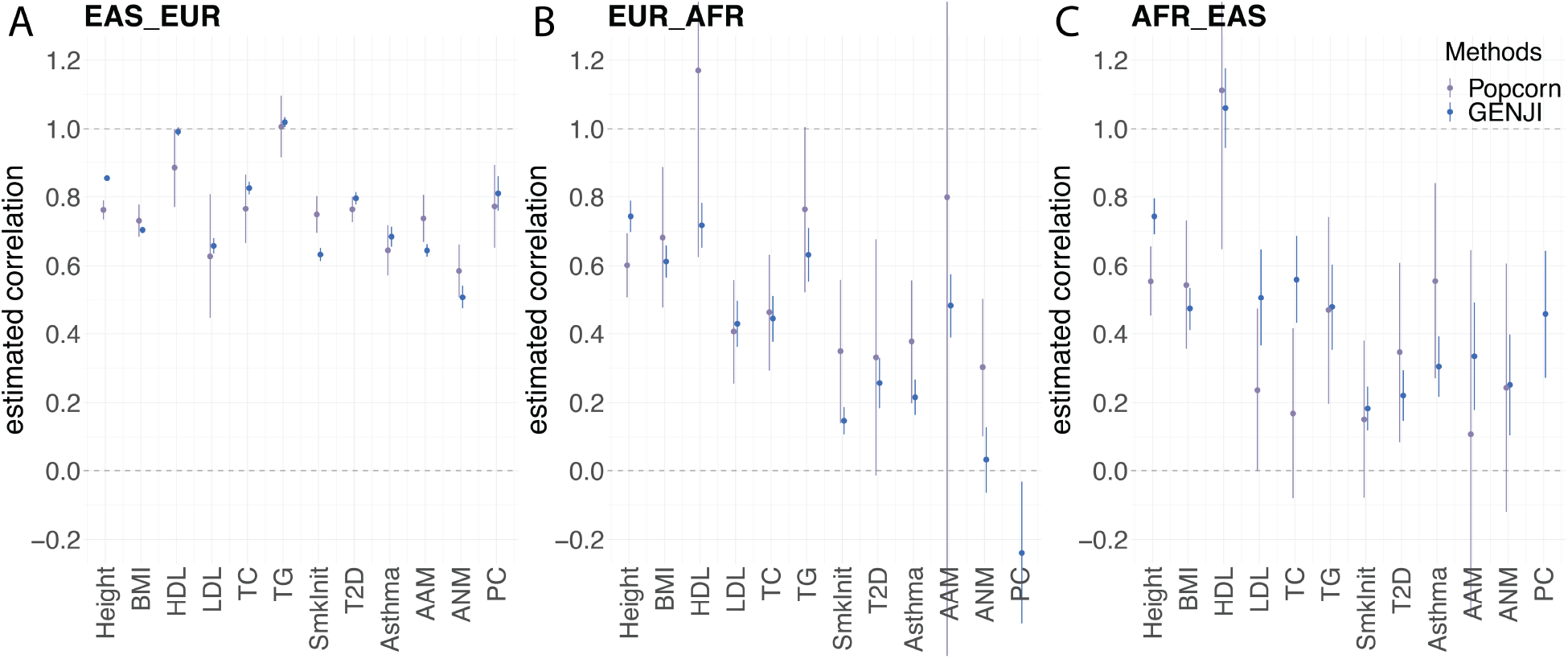
Transethnic genetic correlation estimates of GENJI and Popcorn for 12 traits cross different populations. To make the comparisons fair, we use in-sample reference panel for Popcorn for the population of which GENJI takes the individual-level data as input. African, East Asian and European are denoted as AFR, EAS, EUR, respectively. The point and the point range in the figures demonstrate the point estimates and their standard errors for transethnic genetic correlation of Popcorn and GENJI between **(A)** East Asian and European populations, **(B)** European and African populations, and **(C)** African and East Asian populations. Some of the point ranges are truncated due to their extraordinary values. Some of the estimates of Popcorn are not available due to low heritability estimation. We summarized the abbreviations and detailed information about the data sources in **Supplementary Table 5** and **7**.

### GENJI improves cross-population risk prediction

To illustrate the benefits of improved transethnic genetic correlation estimation, we show that cross-population PRS is more predictive with the help of more accurate genetic correlation estimates. It has been shown that leveraging transethnic genetic correlation can substantially improve PRS performance especially for minority groups^31^. Here, based on a commonly used method for computing PRS, clumping and thresholding (C+T)^32–34^, we highlight the advances of GENJI that the cross-population PRS improved by genetic correlation estimates of GENJI is more predictive than PRSs constructed in other ways.

We constructed the PRSs of Height and BMI for the samples of African ancestry in Population Architecture Genomics and Epidemiology (PAGE): Multiethnic Cohort (MEC) (n=3,520). The genetic weights were trained from UKBB data. We used the estimates of transethnic genetic correlation between African and European populations in the previous section to improve the prediction (**Methods**). We also included PRS constructed by meta-analysis of African and European populations using METAL^35^ in the comparison.

The predictive R^2^ of PRSs improved by GENJI were higher than all the other approaches (**Figure 6**) as a result of the more accurate estimates from GENJI. The R^2^ of the PRSs that only were constructed by GWAS of African population were saliently the lowest. Since the transethnic genetic correlation of BMI is relatively low (GENJI corr=0.61) and the transethnic genetic correlation of BMI estimated by Popcorn (Popcorn corr=0.68) is relatively close to that estimated by GENJI (**Figure 5**), PRSs improved by GENJI and Popcorn outperformed the PRS constructed by METAL for BMI. On the other hand, the transethnic genetic correlation of Height is relatively high (GENJI corr=0.74). In addition, the estimate of Popcorn (Popcorn corr=0.60) is far from that of GENJI and might underestimate the transethnic genetic correlation for Height (**Figure 5**). Consequentially, PRS improved by GENJI and the PRS constructed by METAL outperformed PRS improved by Popcorn for Height.

**Figure 6.**
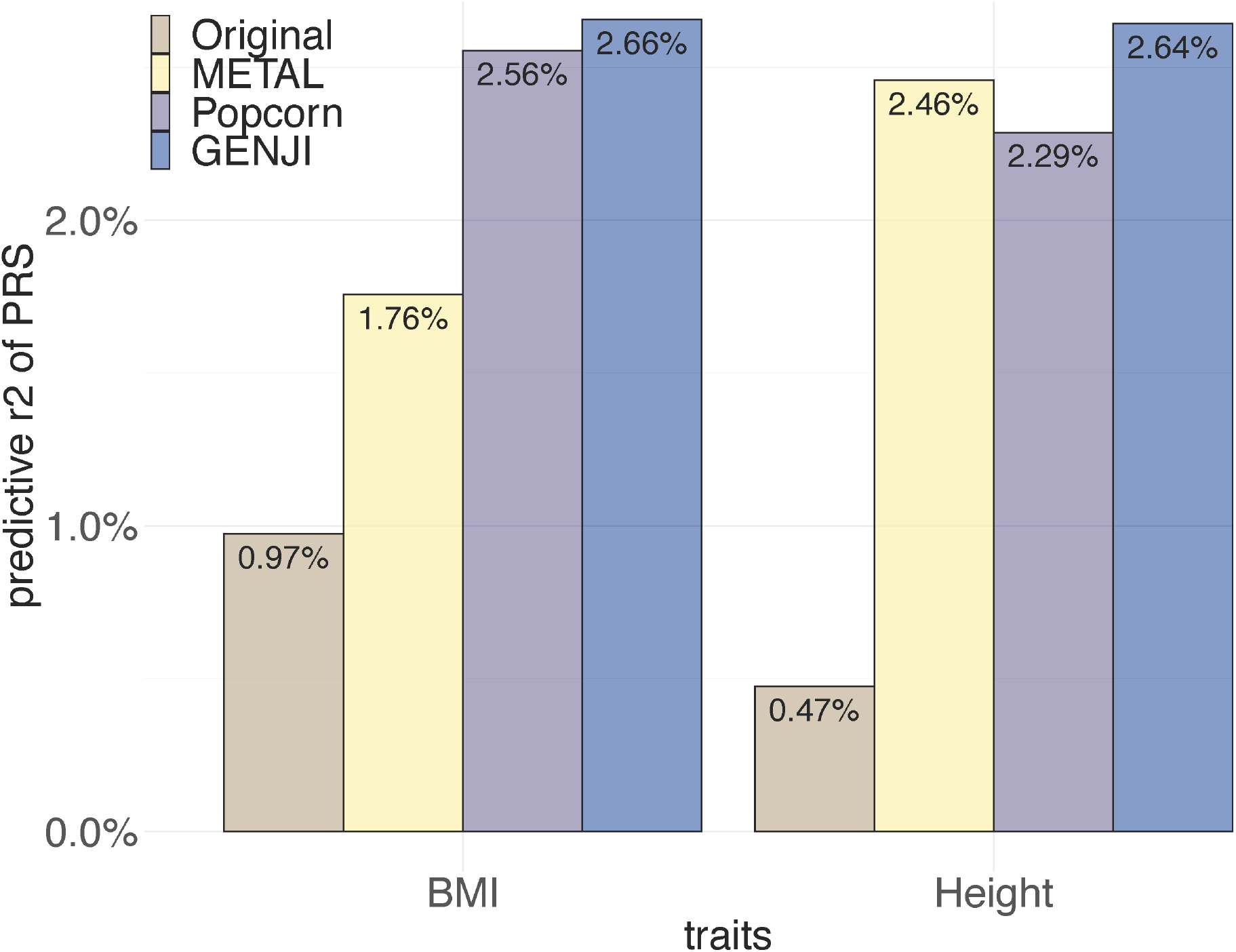
The R^2^ of the cross-population PRS. The R^2^ of the PRSs for BMI and Height are visualized by bar plots. The exact values of R^2^ are annotated in the figure. The PRSs are constructed by four different ways: (1) Original, only using the GWAS of African population to construct the PRS; (2) METAL, using meta-analysis of GWASs of African and European populations to construct the PRS; (3) Popcorn, using estimates of Popcorn to adjust the genetic weights in PRS; (4) GENJI, using estimates of GENJI to adjust the genetic weights in PRS.

## Discussion

Genetic correlation is a powerful metric to effectively measure the etiologic sharing of numerous phenotypes. Recently, increasing attention has been paid to the genetic architecture of non-European populations^36–38^. Transethnic genetic correlation can provide insights to study how the genetic architecture of complex phenotypes varies between populations^14^. Methods to estimate genetic correlation based on individual-level data or summary-level data have achieved some success. The individual-level data based methods (e.g. GREML^7^) have proved to be statistically more efficient than the summary-level data based methods^6^. When individual-level data are available for one trait and summary-level data are available for the other trait, to estimate genetic correlation, researchers have to transform the individual-level data to summary statistics and apply summary-statistics-based methods, leading to information loss and suboptimal estimates. Now that individual-level data are increasingly accessible, we propose a statistical framework that can estimate within-population or transethnic genetic correlation with individual-level data for one trait and summary-level data for the other trait which can provide more accurate estimates than summary-statistics-based methods.

GENJI provides statistically rigorous and computationally efficient inference for both within-population and transethnic genetic correlations and substantially outperforms summary data-based methods in our simulations. We also observe that the information loss from individual-level data to summary-level data is likely to be induced by estimation of covariance of non-genetic effects for overlapped samples which parameterized as the intercept of LDSC. The point estimation of LDSC with intercept fixed at zero is comparable with GREML and GENJI and is more efficient than LDSC without constraint on intercept by 30% under the circumstance of no overlapped samples. Since LDSC uses block jackknife to estimate the standard errors, the statistical power of LDSC is suboptimal no matter whether its intercept is fixed or not. The “samples” in the regression model of GENJI are the individuals in the studies. So, if the GWAS samples are unrelated, we can directly compute the standard errors of the estimates in the close form instead of applying resampling based methods.

Applied to different datasets to estimate within-population genetic correlation, GENJI consistently identified more significant trait pairs than LDSC. The findings of GENJI have been validated by GREML on both point estimation and false discovery control. Notably, GENJI identified 148% more trait pairs than LDSC in an application to 150 trait pairs from UKBB and 25 publicly available GWAS summary datasets.

GENJI can also yield reliable outcomes for transethnic genetic correlation estimation. Estimates of GENJI are more consistent with those of GREML than the estimates from Popcorn. In addition, the standard errors from GENJI are substantially smaller than those from Popcorn. The results show that the genetic effects of East Asian and European are more correlated than African and European or African and East Asian, which might be interpreted by the closer genetic distance between Europeans and Asians.

Furthermore, for most traits, we observe that the transethnic genetic correlations across populations are significantly different from either zero or one. These results suggest that more sophisticated methods are in need to study the genetic architectures for non-European populations. On one hand, it is inappropriate to directly generalize the findings in European population to non-European population^36,37^. On the other hand, much amount of information can be borrowed by cross-population analysis. For example, elaborated modeling of GWAS from multiple populations can improve PRS prediction. Researchers have proposed several methods for cross-population PRS which leverage GWAS data from other populations to boost the performance of PRS prediction on the target population^31,38–40^. Genetic correlation has proved to be a powerful way to improve PRS performance^31,41^. PRS prediction is bound to benefit from more accurate estimation of genetic correlation. As a showcase, we used the estimates for transethnic genetic correlation between African and European populations from GENJI and Popcorn to construct cross-population PRS for African population. We compared the PRS improved by genetic correlation with two alternative approaches: PRS based on the target population alone and PRS built on a meta-analysis combing multiple populations, which are equivalent to the cases of genetic correlation equal to zero and one, respectively. The performance of PRS can be evaluated by predictive R^2^ in external testing datasets. We find that PRS improved by the GENJI outperformed other approaches, which suggests that estimation from GENJI is closer to the underlying true value.

Our method has some limitations. First, to implement the weighted regression in GENJI, if the two studies share non-zero samples, we need to know who is included in the set of the shared samples. Second, the advantage of GENJI diminishes with the increase of the proportion of overlapping samples. Too many overlapping samples between the studies make the regression problem in GENJI suffer from multicollinearity and result in unstable estimates (**Methods**). We note that these two limitations are both induced by sample overlapping, which does not exist in transethnic genetic correlation applications. Third, genome-wide genetic correlations only reflect the average concordance of genetic effects across the genome and often fail to reveal the local, heterogenous pleiotropic effects, especially when the underlying genetic basis involves multiple etiologic pathways^42–44^. Future directions include extending our method to estimate local genetic correlation jointly using individual-level and summary-level data.

Taken together, GENJI provides a biologically-motivated and statistically principled analytical strategy to tackle etiologic sharing of complex traits within and across populations. A combination of individual-level and summary-level data is more desired to fully utilize the information provided by the data. Moreover, it is doable in prospect owing to increasing availability of individual-level data. We believe GENJI will have wide applications in complex traits and cross-population genetics researches.

## Methods

### Statistical model

The statistical frameworks of GENJI for the estimations of within-population genetic correlation and transethnic genetic correlation are nearly the same. Assume that the two studies have sample size *n*_1_ and *n*_2_, respectively. Standardized trait values *ϕ*_1_ and *ϕ*_2_ follow the linear models below:

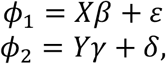

where *X* and *Y* are *n*_1_ × *m* and *n*_2_ × *m* standardized genotype matrices; *m* is the number of shared SNPs between the two studies; *ε* and *δ* are the noise terms; and *β* and *γ* denote the genetic effects for *ϕ*_1_ and *ϕ*_2_. The genetic covariance *ρ_g_* is defined as the covariance of the (population-specific) allele-variance-normalized random SNP effect sizes. So, the combined random vector of *β* and *γ* follows a multivariate normal distribution given by:

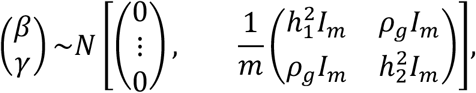

where *I_m_* is the identity matrix of size *m*. In the estimation of transethnic genetic correlation, this assumption for the distribution of SNP effect sizes corresponds to genetic-impact correlation in Popcorn paper^14^. Since trait values *ϕ*_1_ and *ϕ*_2_ are standardized, we have 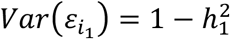 and 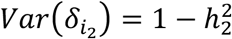 for 1 ≤ *i*_1_ ≤ *n*_1_ and 1 ≤ *i*_2_ ≤ *n*_2_. The covariance of genetic factors of the two traits is equal to *ρ_g_*. In fact, for an individual with normalized genotype vector *X*_0_, 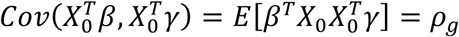. Here we assume that we have individual-level data for the first study and only summary-level data for the second study. With this assumption, we treat *X* as a fixed genotype matrix, and *Y* as an unknown random design matrix. We further assume that each row of *Y* is independently drawn from a distribution with covariance matrix *R*, the *m* × *m* LD matrix for study 2, i.e., *Cov*(*Y*_*i*_2__.) = *R*. The LD matrix can be estimated from external reference panel (e.g., 1000 Genomes Project^27^) or the genotype of study 1 if the two studies are conducted on the same population. For the phenotype vectors *ϕ*_1_ and *ϕ*_2_, we assume *ϕ*_1_ is known while *ϕ*_2_ is unknown. For GWAS summary data of study 2, we only observe z-scores and we can approximate z-score of SNP *j* by 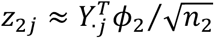. We use *z*_2_ to denote the vector of length *m* which contains all the z-scores in study 2.

In practice, for within-population genetic correlation estimation, two different studies may share a subset of samples. Without loss of generality, we assume the first *n_s_* samples in each study are shared (*n_s_* ≤ *n*_1_ and *n_s_* ≤ *n*_2_). The non-genetic effects of the shared samples for the two studies are correlated:

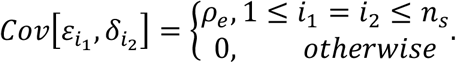

We denote the genotype of the overlapped samples as *X_s_*, which consists of the first *n_s_* rows of *X*. We note that the genotype matrix *X_s_* is known to us because *X_s_* is a submatrix of *X*. We also note that the phenotypes for study 2 of the overlapped samples are unknown although we know their genotypes.

### Parameter estimation

The primary parameter to be estimated is genetic covariance *ρ_g_*. To fully utilize the individual-level information, we relate the expectation of 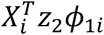 to the parameter of interest, where *X_i_* and *ϕ*_1*i*_ are the genotype vector and phenotype value of the *i*th sample in study 1, respectively. It can be shown that

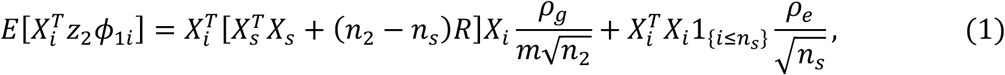

where 1_{*i*≤*n_s_*}_ is the indicator function for the overlapped samples. We estimate genetic covariance by regressing 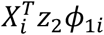 against 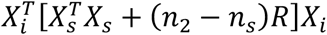 and then multiplying the resulting slope by 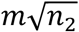. We note that for transethnic genetic covariance, since there is no overlapp sample between the studies, (1) reduces to

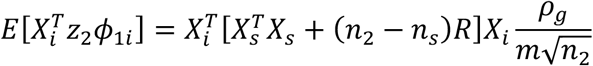

and we constrain the intercept of the regression to be zero.

To obtain the optimal estimator of the regression, we apply weighted regression. The weights are given by the reciprocal of the variance of 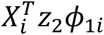:

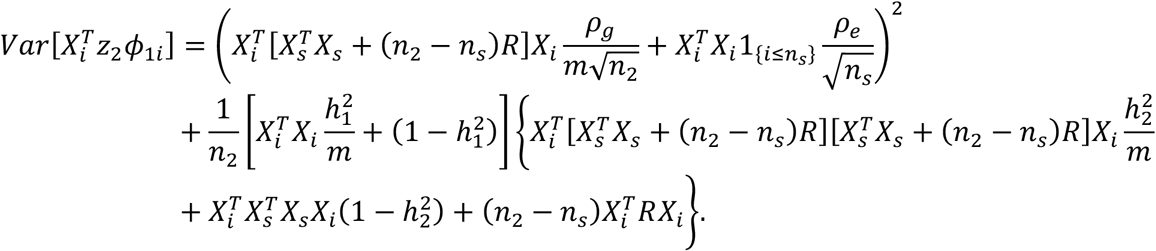

The weights depend on 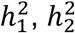, *ρ_g_* and *ρ_e_* that are unknown. The parameters for heritability 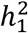 and 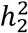 can be estimated by GREML and LDSC, respectively. For *ρ_g_* and *ρ_e_*, we estimate them in two steps. In the first step, we set *ρ_g_* = *ρ_e_* = 0 to calculate the weights. Then, we use the estimates of *ρ_g_* and *ρ_e_* in the first step to calculate the weights for the second step. The result of genetic covariance estimation is the estimate from the second regression.

The estimate for genetic covariance is on the observed scale if one or both studies are case-control study. The observed scale genetic covariance is 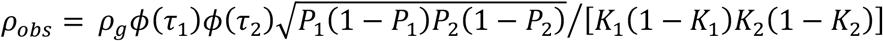 when both studies are case-control studies, where *ρ_g_* is the liability scale genetic covariance, *ϕ* is the standard normal density, *τ*_1_ and *τ*_2_, *P*_1_ and *P*_2_, and *K*_1_ and *K*_2_ are the liability threshold, sample prevalence, and population prevalence of study 1 and study 2, respectively. If one study is case-control study, the observed scale genetic covariance is given by 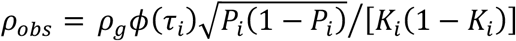, *i* = 1,2. The detailed derivations are similar to those presented in the Supplementary Note of Zhang et al.^42^.

Since the samples are independent to each other, we can directly use the standard errors of the coefficients estimated by the weighted regression. Z-test is applied to determine the statistical significance. For transethnic genetic correlation, we also use the standard errors from weighted regression to test whether the genetic correlation is significantly below 1 with one-tailed test.

We can extend the above method to include the covariates of study 1:

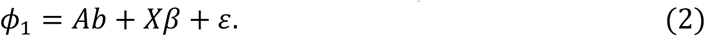

Here *A* is an *n*_1_ × *p* matrix of covariates and *b* is a vector of length *p* for fixed effects. Let *p*_0_ be the rank of *A* and *U* be an *n*_1_ × (*n*_1_ – *p*_0_) matrix that consists of the orthonormal bases of the orthogonal complement space of *A. U* satisfies *U^T^U* = *I*_*n*_1_−*p*_0__ and 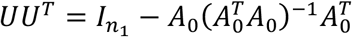, where *A*_0_ is an *n*_1_ × *p*_0_ full rank submatrix of *A*. We multiply by *U^T^* on both sides of (2) which yields:

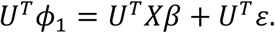

We can simply use standardized *U^T^ϕ*_1_ as the corrected phenotypes and column standardized *U^T^X* as the corrected genotypes and then input the corrected phenotypes and genotypes to the regression introduced above in (1).

### Simulation settings for within-population genetic correlation

In this section, we introduce the procedure that we use simulated data to compare the performance of GENJI, LDSC, and GREML. Phenotypes in our simulations were generated based on unimputed genotype data from the WTCCC for within-population genetic correlation estimation. Samples were randomly divided into two equal subgroups and each subgroup had 7,959 individuals. We denote them as set 1 and set respectively. We randomly sampled individuals from set 1 and set 2 and created set set 4 and set 5 whose sample sizes were all equal to 7,959 such that they had 10%, 25% and 50% overlapped samples with set 1, respectively. Genotype data of 503 individuals with European ancestry from the 1000 Genomes Project phase III^27^ were used as the LD reference in our simulations. SNPs with ambiguous alleles or minor allele frequencies (MAF) less than 5% were removed. 253,196 SNPs presented in both WTCCC and 1000 Genomes Project remained.

The effect sizes of SNPs were generated by a multivariate normal distribution and we applied Genome-wide Complex Trait Analysis (GCTA)^9^ to simulate the phenotypes. We used PLINK^34^ to run GWAS and obtain summary statistics of the two simulated phenotypes. We repeated each simulation setting 100 times. Detailed simulation settings are summarized below.

We fixed the heritability of the phenotypes as 0.5 and the values of genetic covariance were set from 0 to 0.25. The first GWAS was simulated on the individuals in set 1 and the second GWAS was simulated for set 2, set 3, set 4 and set 5 in four scenarios with different sample overlapping. The covariance of non-genetic effects on overlapped samples was set to be 0.2, i.e., *ρ_e_* = 0.2. When there was no overlapped sample between the two studies, we also included LDSC with intercept fixed at zero in our comparison. GCTA was applied for GREML implementation.

We also simulated binary traits using the liability model for no-sample-overlap scenario to investigate the performance of the methods for case-control study. The liability scale genetic covariance was still set from 0 to 0.25. We set the threshold for the liability to be 80% quantile of standard normal distribution such that the prevalence of the binary traits was 0.2.

### Simulation settings for transethnic genetic correlation

In this section, we used similar procedure introduced above to compare the performance of GENJI, Popcorn, and GREML on transethnic genetic correlation estimation. The unimputed European genotype data from the WTCCC (set 1), and genotype data of genetically unrelated UKBB African ancestry samples were used to generate phenotypes, with sample sizes to be 7,959 and 6,992, respectively. We provided GENJI with individual-level data of African ancestry from UKBB and summary-level data generated from WTCCC European samples. Individuals of European and African ancestry from the 1000 Genomes Project phase III were used as the reference for Popcorn. For fair comparison, we also added an additional setting for Popcorn, where the reference panel was the genotype data used to generate the phenotype of the African traits (in-sample reference) because GENJI uses the individual-level data of the GWASs of African ancestry to estimate genetic correlation. SNPs with ambiguous alleles or MAFs less than 5% were removed. 159,093 shared SNPs remained in genotype data of WTCCC, UKBB, and 1000 Genomes Project.

Genetic effect sizes were generated by multivariate normal distribution. Similarly, phenotypes and summary statistics were generated by GCTA and PLINK, respectively. The heritability of the phenotypes was still set to be 0.5, and the values of genetic covariance were set to range from 0 to 0.5. Each setting was repeated 100 times. We used the genetic impact correlation for Popcorn. Since it was also of great interest to investigate whether the genetic effects of a specific trait between two populations were perfectly correlated (transethnic genetic correlation=1), besides the traditional null that genetic correlation is equal to zero, we also tested for *H*_0_: *corr* = 1 with alternative *H*_1_ = *corr* < 1.

### Within-population genetic correlation estimation for European ancestry GWASs

We used GREML, LDSC, and GENJI to estimate genetic correlations across the 12 phenotypes in WTCCC and NFBC datasets. We downloaded genotype and phenotype data of NFBC from dbGaP^45^ (accession: phs000276.v2.p1). Since only a small proportion of SNPs were shared by these these two datasets, we imputed WTCCC using the Michigan Imputation server^46^ which ended up with 227,383 shared SNPs with NFBC with MAF larger than 0.05. Sex, age, and top 4 principal components were included as covariates to implement GREML, GENJI and perform GWAS. To run GENJI and LDSC, we generated summary data for both cohorts using PLINK. We provided GENJI with individual-level data from NFBC and summary data from WTCCC. Genotypes of European ancestry in 1000 Genomes Project were still used as reference panel for GENJI and LDSC. The details of the phenotypes and samples sizes are summarized in **Supplementary Tables 1** and **2**.

We then applied GENJI on GWASs of European ancestry with much larger sample sizes. We estimated the genetic correlation between 25 common traits with publicly available GWAS summary data and 6 common traits from UKBB. We included the genotypes of 276,731 genetically unrelated samples of European ancestry and 306,579 Axiom Array (unimputed) SNPs with MAF larger than 0.05 from UKBB in the analysis. All the trait pairs of WTCCC and NFBC were also included between UKBB traits and the 25 GWASs summary data. Due to the unavailability of the individual-level data of 25 GWASs, GREML was not implemented. Sex, age, and top 4 principal components of UKBB samples were included as covariates to implement GENJI and perform GWAS. We transformed GWASs of UKBB traits to summary data by PLINK as the input to LDSC. Since there was no overlapped sample between UKBB and the 25 GWASs, we fixed the intercept of LDSC at zero. The details about the phenotypes and samples sizes of the UKBB traits are given in **Supplementary Table 5**. The details about the sample sizes and the sources of the 25 GWASs are given in **Supplementary Table 4**.

### Transethnic genetic correlation estimation for European, African, and East Asian ancestry GWASs

We used individual-level data of samples with European ancestry and African ancestry in UKBB and summary-level data from BBJ^17,18^ to estimate transethnic genetic correlation of 12 traits across the three populations. Our choice of these traits was based on the availability of the data and the heritability of the traits. All the chosen traits have nominally significant heritability (p<0.05) across all the populations (**Supplementary Table 12**). We used GWAS summary data from BBJ because they are publicly available and have larger sample sizes than samples of East Asian ancestry in UKBB. Similarly, we only used unimputed SNPs from UKBB. We used the genetic impact correlation for Popcorn. Detailed information about datasets used in this section is summarized in **Supplementary Tables 5** and **7**.

We first compared the performance of GENJI with Popcorn by the consistency with GREML on transethnic genetic correlation estimation between the European ancestry and African ancestry populations. Since it is of expensive computational burden to implement GREML on Biobank-scale data (n=276,731 for samples of European ancestry), we randomly extracted 10,000 individuals of European ancestry and performed the three methods on the GWASs based on this subset. Sex, age, and top 4 principal components were included as covariates. We provided GENJI with individual-level data of African ancestry and summary-level generated from European ancestry. For African population, Genotype data of African ancestry from 1000 Genome Project (external reference panel) and UKBB (in-sample reference panel) were used as reference panel to estimate LD in two implementations of Popcorn, respectively, for the sake of comparison fairness between GENJI and Popcorn. For European population, genotype data of European ancestry in 1000 Genomes Project (external reference panel) was used as reference panel for both GENJI and Popcorn.

Then, GENJI and Popcorn were further compared on the estimation of transethnic genetic correlation among European, African, and East Asian ancestry GWASs with larger sample sizes. For the results between African and European populations, genetic correlations were estimated using the whole dataset of UKBB. Similarly, we provided GENJI with individual-level data of African ancestry and summary-level generated from European ancestry. Individuals of African ancestry from UKBB (in-sample reference panel) and of European ancestry from the 1000 Genomes Project were used as the reference for Popcorn. GENJI also used European ancestry genotype from the 1000 Genomes Project as the reference panel. The GWAS summary data for East Asian population were downloaded from BBJ website (**URLs**). For the genetic correlation between African and East Asian populations, and European and East Asian populations, we provided GENJI with individual-level data from UKBB (African or European population) and summary data for East Asian population. Individuals of African or European ancestry in UKBB (in-sample reference panel), and of East Asian ancestry from the 1000 Genomes Project were used as the reference for Popcorn while GENJI used the genotype of East Asian ancestry from 1000 Genomes Project as its reference panel.

### Using transethnic genetic correlation to improve cross-population PRS

We used the transethnic genetic correlation estimated by GENJI and Popcorn to improve PRS prediction for Height and BMI on African population. We compared the PRS adjusted by genetic correlation with the unadjusted PRS and the PRS based on the meta-analysis combining the GWASs of European and African populations using METAL^35^. We used the GWAS data of African and European populations for Height and BMI from UKBB as the training data. The transethnic genetic correlations for Height and BMI were directly applied from the previous section. Samples of African ancestry from PAGE: MEC were included in the testing set. We downloaded the individual-level phenotype and genotype data from dbGaP^45^ (accession: phs000220.v2.p2). We imputed genotype from PAGE using the Michigan Imputation server^46^. We restricted the analysis to autosomal variants with genotype missing rate per marker < 0.05, missing rate per individual < 0.1, Hardy-Weinberg Equilibrium p-value > 1e-9, and MAF > 0.05. After quality control, we ended with 3,520 samples of African ancestry with both genotype and phenotype data available. 1,334,601 SNPs remained in the imputed genotype data of PAGE remain.

For the unadjusted PRS, we first clumpedd the SNPs by PLINK^34^. We set the significance threshold for index SNPs as 1, LD threshold for clumping as 0.1, and physical distance threshold for clumping as 250 kb. We then used --*score* function in PLINK to calculate the unadjusted PRS. The PRS from meta-analysis was constructed similarly. We used the inverse variance based analytical strategy given by Table 1 of METAL^35^ and generate genetic weights to compute PRS.

For PRS adjusted by genetic correlation, we applied similar methods in Multi-Trait Analysis of GWAS (MTAG)^47^ and Multi-Ancestry Meta-Analysis (MAMA)^48^ to adjust the effect sizes of SNPs. The effect size vector for the first study (population) is *β* which is used to construct the PRS for the target population. The effect size vector for the second study (population) is *γ* which is the auxiliary data. We only used the SNPs after clumping and hence the SNPs were considered to be independent (no LD). So, without LD, conditioned on the true effect size *β_i_*, the marginal effect of normalized genotype for SNP *i* from the GWAS of the target population is subject to normal distribution: 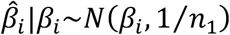. The conditional distribution of 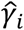 given *β_i_* is:

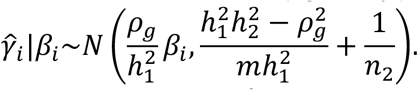

Finally, the conditional expectation of *β_i_* given 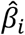 and 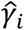 is:

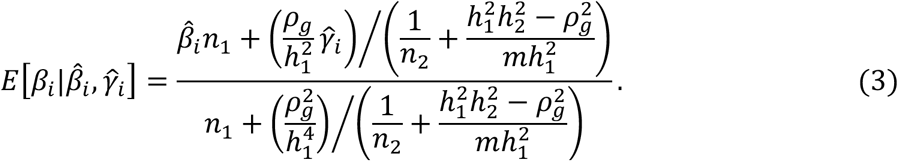

We plugged the estimates from GENJI and Popcorn into (3) and used the results given by (3) as the genetic weight improved by transethnic genetic correlation to compute the new PRS.

## URLs

GCTA (https://cnsgenomics.com/software/gcta/#GREML)

LDSC (https://github.com/bulik/ldsc)

Popcorn (https://github.com/brielin/popcorn)

UKBB (https://www.ukbiobank.ac.uk)

WTCCC (https://www.wtccc.org.uk)

BBJ (http://ienger.riken.jp/)

PLINK (https://zzz.bwh.harvard.edu/plink/profile.shtml)

Michigan Imputation Server (https://imputationserver.sph.umich.edu/index.html#!)

## Data and code availability

GENJI software is publicly available at https://github.com/YiliangTracyZhang/GENJI

## Acknowledgements

This study was Supported in part by NIH grants 3P30AG021342-16S2 and 1R01GM122078 and NSF grants DMS 1713120 and DMS 1902903.

This study makes use of data generated by the Wellcome Trust Case-Control Consortium. We conducted the research using the UKBB resource under approved data requests (refs: 29900). This study makes use of summary statistics from many GWAS consortia. We thank the investigators in these GWAS consortia for generously sharing their data. The NFBC1966 Study is conducted and supported by the National Heart, Lung, and Blood Institute (NHLBI) in collaboration with the Broad Institute, UCLA, University of Oulu, and the National Institute for Health and Welfare in Finland. This manuscript was not prepared in collaboration with investigators of the NFBC1966 Study and does not necessarily reflect the opinions or views of the NFBC1966 Study Investigators, Broad Institute, UCLA, University of Oulu, National Institute for Health and Welfare in Finland and the NHLBI. Funding support for the PAGE Multiethnic Cohort study was provided through the National Cancer Institute (R37CA54281, R01CA6364, P01CA33619, U01CA136792, and U01CA98758) and the National Human Genome Research Institute (U01HG004802). Assistance with phenotype harmonization, SNP selection, data cleaning, meta-analyses, data management and dissemination, and general study coordination, was provided by the PAGE Coordinating Center (U01HG004801-01). GWAS data of east Asian ancestry were downloaded from BBJ website, which were all publicly available.

## Author contributions

Y.Z. conceived and designed the study.

Y.Z. developed the statistical framework.

Y.Z. implemented the software.

Y.Z. and Y.C. performed simulations.

Y.Z., Y.C., and Y.Y. performed real data analysis.

Y.Y. processed UKBB data.

W.J downloaded and curated NFBC and PAGE data.

Y.Y, Q.L., and H.Z. advised on the biological interpretation of the real data findings.

W.J., Q.L., and H.Z. advised on statistical and genetics issues.

Y.Z. and Y.C. wrote the manuscript.

All authors contributed in manuscript editing and approved the manuscript.

## References

1. Visscher, P.M. et al. 10 Years of GWAS Discovery: Biology, Function, and Translation. American Journal of Human Genetics 101, 5–22 (2017).

2. Lichtenstein, P. et al. Common genetic determinants of schizophrenia and bipolar disorder in Swedish families: a population-based study. Lancet 373, 234–9 (2009).

3. Grove, J. et al. Identification of common genetic risk variants for autism spectrum disorder. Nat Genet 51, 431–444 (2019).

4. Anttila, V. et al. Analysis of shared heritability in common disorders of the brain. Science 360, 1313-+ (2018).

5. Cross-Disorder Group of the Psychiatric Genomics Consortium. Electronic address, p.m.h.e. & Cross-Disorder Group of the Psychiatric Genomics, C. Genomic Relationships, Novel Loci, and Pleiotropic Mechanisms across Eight Psychiatric Disorders. Cell 179, 1469–1482 e11 (2019).

6. Zhang, Y. et al. Comparison of methods for estimating genetic correlation between complex traits using GWAS summary statistics. Briefings in Bioinformatics (2021).

7. Lee, S.H., Yang, J., Goddard, M.E., Visscher, P.M. & Wray, N.R. Estimation of pleiotropy between complex diseases using single-nucleotide polymorphism-derived genomic relationships and restricted maximum likelihood. Bioinformatics 28, 2540–2542 (2012).

8. Bulik-Sullivan, B. et al. An atlas of genetic correlations across human diseases and traits. Nature Genetics 47, 1236-+ (2015).

9. Yang, J., Lee, S.H., Goddard, M.E. & Visscher, P.M. GCTA: a tool for genome-wide complex trait analysis. Am J Hum Genet 88, 76–82 (2011).

10. Lee, S.H. & van der Werf, J.H. MTG2: an efficient algorithm for multivariate linear mixed model analysis based on genomic information. Bioinformatics 32, 1420–2 (2016).

11. Wu, Y. et al. Fast estimation of genetic correlation for Biobank-scale data. bioRxiv, 525055 (2020).

12. Zheng, J. et al. LD Hub: a centralized database and web interface to perform LD score regression that maximizes the potential of summary level GWAS data for SNP heritability and genetic correlation analysis. Bioinformatics 33, 272–279 (2017).

13. Ni, G., Moser, G., Schizophrenia Working Group of the Psychiatric Genomics, C., Wray, N.R. & Lee, S.H. Estimation of Genetic Correlation via Linkage Disequilibrium Score Regression and Genomic Restricted Maximum Likelihood. Am J Hum Genet 102, 1185–1194 (2018).

14. Brown, B.C., Ye, C.J., Price, A.L., Zaitlen, N. & Network, A.G.E. Transethnic Genetic-Correlation Estimates from Summary Statistics. American Journal of Human Genetics 99, 76–88 (2016).

15. Sudlow, C. et al. UK biobank: an open access resource for identifying the causes of a wide range of complex diseases of middle and old age. PLoS Med 12, e1001779 (2015).

16. Wellcome Trust Case Control, C. Genome-wide association study of 14,000 cases of seven common diseases and 3,000 shared controls. Nature 447, 661–78 (2007).

17. Kanai, M. et al. Genetic analysis of quantitative traits in the Japanese population links cell types to complex human diseases. Nat Genet 50, 390–400 (2018).

18. Ishigaki, K. et al. Large-scale genome-wide association study in a Japanese population identifies novel susceptibility loci across different diseases. Nat Genet 52, 669–679 (2020).

19. Scott, R.A. et al. An Expanded Genome-Wide Association Study of Type 2 Diabetes in Europeans. Diabetes 66, 2888–2902 (2017).

20. Yengo, L. et al. Meta-analysis of genome-wide association studies for height and body mass index in approximately 700000 individuals of European ancestry. Hum Mol Genet 27, 3641–3649 (2018).

21. Teslovich, T.M. et al. Biological, clinical and population relevance of 95 loci for blood lipids. Nature 466, 707–13 (2010).

22. American Diabetes, A. 8. Obesity Management for the Treatment of Type 2 Diabetes: Standards of Medical Care in Diabetes-2020. Diabetes Care 43, S89–S97 (2020).

23. Emerging Risk Factors, C. et al. Major lipids, apolipoproteins, and risk of vascular disease. JAMA 302, 1993–2000 (2009).

24. Oda, E. & Kawai, R. High-density lipoprotein cholesterol is positively associated with hypertension in apparently healthy Japanese men and women. Br J Biomed Sci 68, 29–33 (2011).

25. Bønaa, K. & Thelle, D. Association between blood pressure and serum lipids in a population. The Tromsø Study. Circulation 83, 1305–1314 (1991).

26. Shimizu, Y. et al. Association between high-density lipoprotein-cholesterol and hypertension in relation to circulating CD34-positive cell levels. J Physiol Anthropol 36, 26 (2017).

27. Altshuler, D.M. et al. An integrated map of genetic variation from 1,092 human genomes. Nature 491, 56–65 (2012).

28. Schumacher, F.R. et al. Association analyses of more than 140,000 men identify 63 new prostate cancer susceptibility loci. Nat Genet 50, 928–936 (2018).

29. Day, F.R. et al. Large-scale genomic analyses link reproductive aging to hypothalamic signaling, breast cancer susceptibility and BRCA1-mediated DNA repair. Nat Genet 47, 1294–1303 (2015).

30. Conti, D.V. et al. Publisher Correction: Trans-ancestry genome-wide association meta-analysis of prostate cancer identifies new susceptibility loci and informs genetic risk prediction. Nat Genet 53, 413 (2021).

31. Cai, M. et al. A unified framework for cross-population trait prediction by leveraging the genetic correlation of polygenic traits. Am J Hum Genet 108, 632–655 (2021).

32. Dudbridge, F. Power and predictive accuracy of polygenic risk scores. PLoS Genet 9, e1003348 (2013).

33. International Schizophrenia, C. et al. Common polygenic variation contributes to risk of schizophrenia and bipolar disorder. Nature 460, 748–52 (2009).

34. Purcell, S. et al. PLINK: A tool set for whole-genome association and population-based linkage analyses. American Journal of Human Genetics 81, 559–575 (2007).

35. Willer, C.J., Li, Y. & Abecasis, G.R. METAL: fast and efficient meta-analysis of genomewide association scans. Bioinformatics 26, 2190–1 (2010).

36. Duncan, L. et al. Analysis of polygenic risk score usage and performance in diverse human populations. Nat Commun 10, 3328 (2019).

37. Martin, A.R. et al. Current clinical use of polygenic scores will risk exacerbating health disparities. Nature genetics 51, 584 (2019).

38. Amariuta, T. et al. Improving the trans-ancestry portability of polygenic risk scores by prioritizing variants in predicted cell-type-specific regulatory elements. Nat Genet 52, 1346–1354 (2020).

39. Coram, M.A., Fang, H., Candille, S.I., Assimes, T.L. & Tang, H. Leveraging Multi-ethnic Evidence for Risk Assessment of Quantitative Traits in Minority Populations. Am J Hum Genet 101, 638 (2017).

40. Márquez-Luna, C., Loh, P.R., Consortium, S.A.T.D., Consortium, S.T.D. & Price, A.L. Multiethnic polygenic risk scores improve risk prediction in diverse populations. Genetic epidemiology 41, 811–823 (2017).

41. Hu, Y.M. et al. Joint modeling of genetically correlated diseases and functional annotations increases accuracy of polygenic risk prediction. Plos Genetics 13 (2017).

42. Zhang, Y. et al. Local genetic correlation analysis reveals heterogeneous etiologic sharing of complex traits. bioRxiv, 2020.05.08.084475 (2020).

43. Shi, H.W.B., Mancuso, N., Spendlove, S. & Pasaniuc, B. Local Genetic Correlation Gives Insights into the Shared Genetic Architecture of Complex Traits. American Journal of Human Genetics 101, 737–751 (2017).

44. Guo, H., Li, J.J., Lu, Q. & Hou, L. Detecting Local Genetic Correlations with Scan Statistics. bioRxiv, 808519 (2019).

45. Mailman, M.D. et al. The NCBI dbGaP database of genotypes and phenotypes. Nat Genet 39, 1181–6 (2007).

46. Das, S. et al. Next-generation genotype imputation service and methods. Nat Genet 48, 1284–1287 (2016).

47. Turley, P. et al. Multi-trait analysis of genome-wide association summary statistics using MTAG. Nature Genetics 50, 229-+ (2018).

48. Turley, P. et al. Multi-Ancestry Meta-Analysis yields novel genetic discoveries and ancestry-specific associations. bioRxiv, 2021.04.23.441003 (2021).

