## Supplementary Figure for "Estimating genetic correlation jointly using individual-level and summary-level GWAS data"

### Supplementary Figures

**Supplementary Figure 1. Evaluation of type I error control among within-population genetic correlation (covariance) estimation methods for GWAS datasets without overlapping sample.** LDSC with intercept fixed at zero is denoted as “LDSC_no_intercept”. **(A)** Type I error when the true genetic correlation is 0. **(B)** qq-plot of p value when the true genetic correlation is set to be 0.

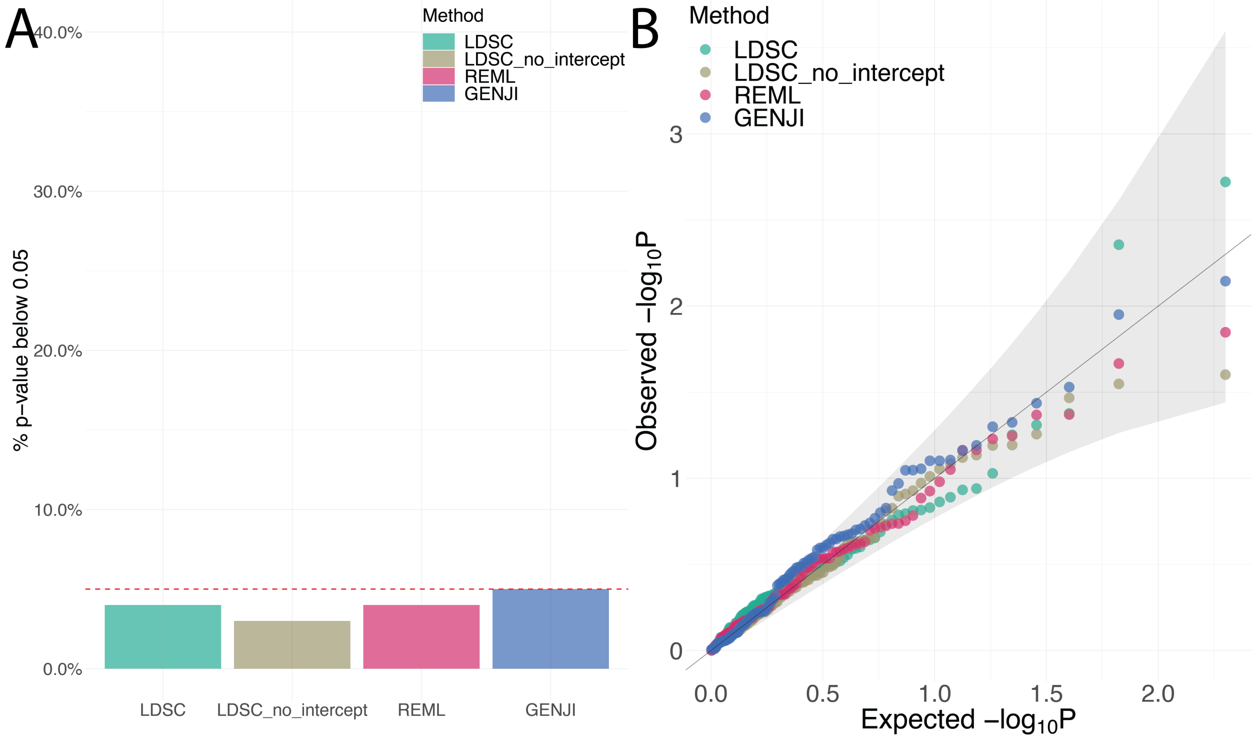

**Supplementary Figure 2. Evaluation of type I error control among within-population genetic correlation (covariance) estimation methods for GWAS datasets with 10% overlapping sample. (A)** Type I error when the true genetic correlation is 0. **(B)** qq-plot of p value when the true genetic correlation is set to be 0.

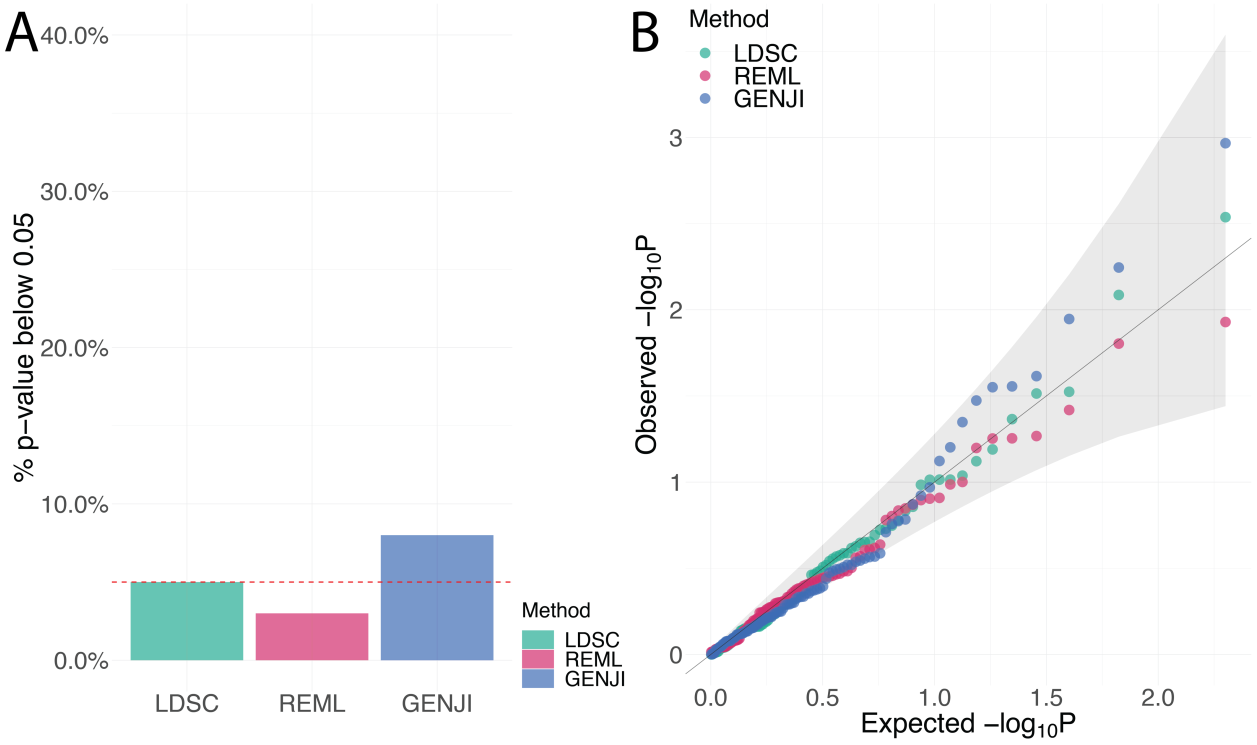

**Supplementary Figure 3. Evaluation of type I error control among within-population genetic correlation (covariance) estimation methods for GWAS datasets with 25% overlapping sample. (A)** Type I error when the true genetic correlation is 0. **(B)** qq-plot of p value when the true genetic correlation is set to be 0.

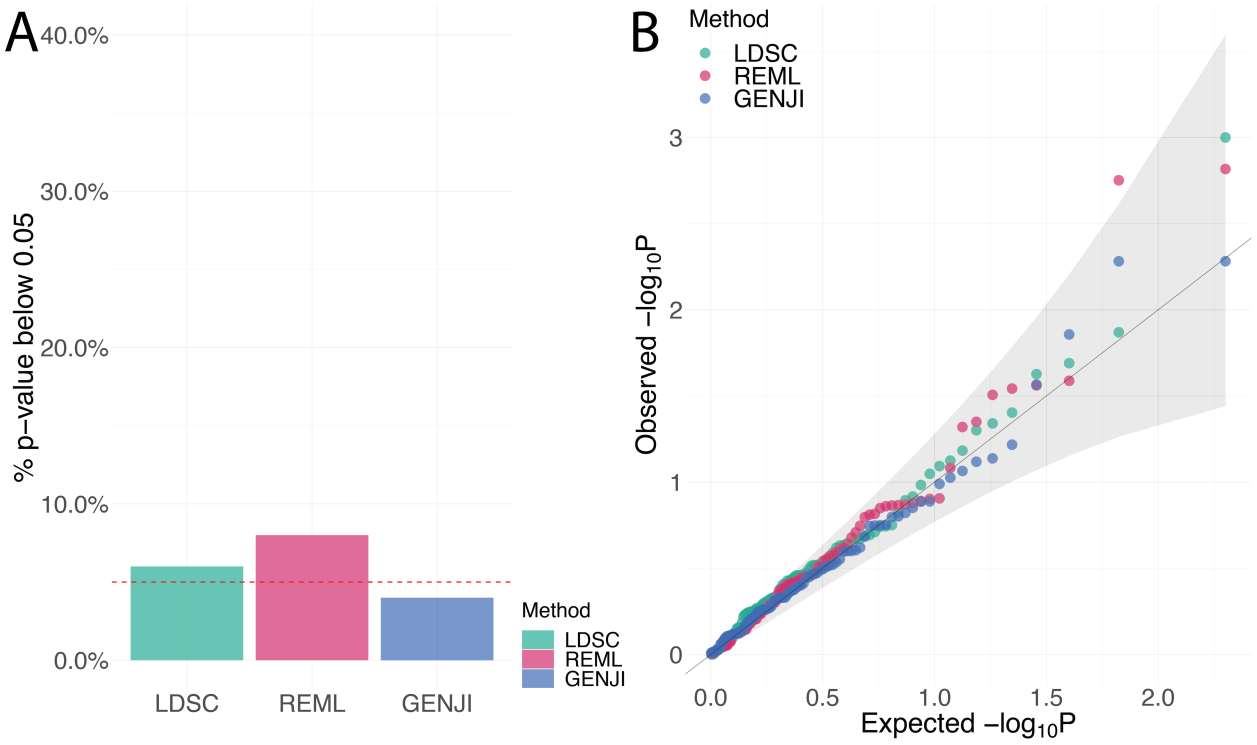

**Supplementary Figure 4. Evaluation of type I error control among within-population genetic correlation (covariance) estimation methods for GWAS datasets with 50% overlapping sample. (A)** Type I error when the true genetic correlation is 0. **(B)** qq-plot of p value when the true genetic correlation is set to be 0.

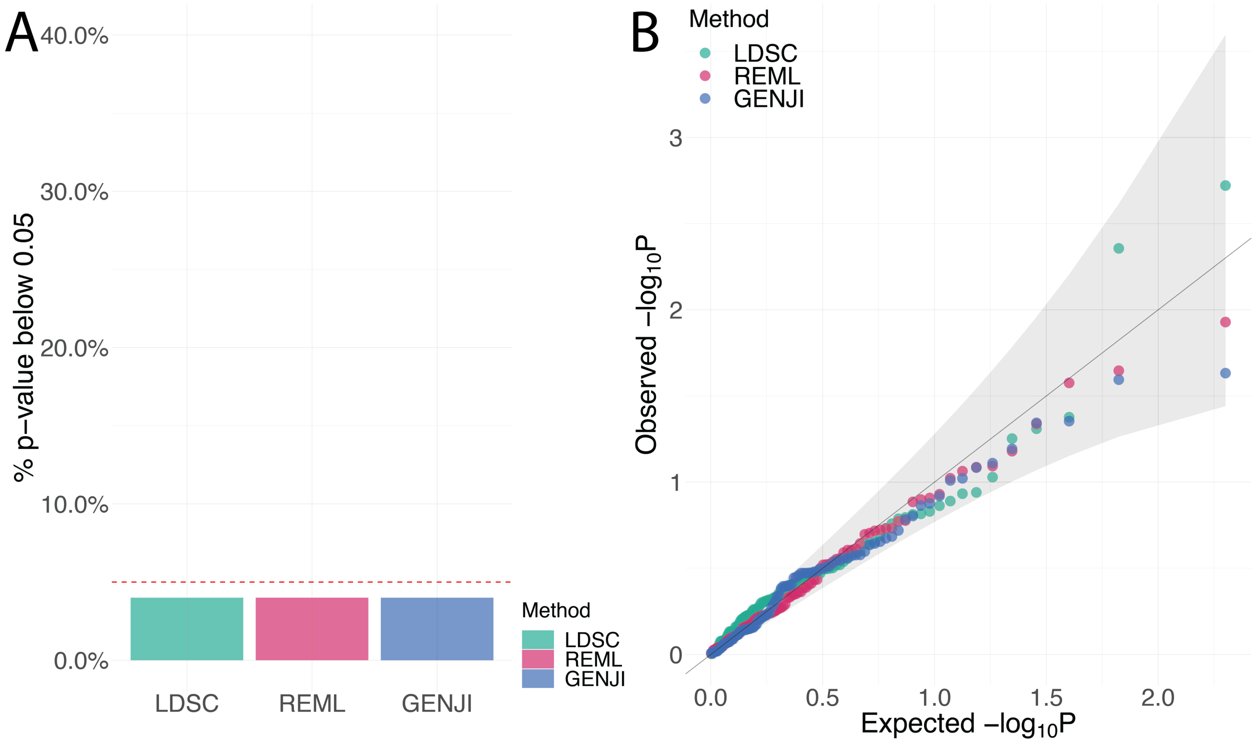

**Supplementary Figure 5. Evaluation of point estimates, type I error control and statistical power among within-population genetic correlation (covariance) estimation methods on binary traits (case-control studies).** The threshold in the liability model to simulate binary traits was set to be $\Phi^{-1}(0.8) = 0.84$. LDSC with intercept fixed at zero is denoted as “LDSC_no_intercept”. **(A)** Genetic correlation among REML, LDSC, and GENJI are demonstrated by boxplot which shows the quantiles of the estimates. The red dashed lines represent true values. **(B)** Type I error and statistical power when true parameters are zero or nonzero, respectively. **(C)** Type I error when the true genetic correlation is 0. **(D)** qq-plot of p value when the true genetic correlation is set to be 0.

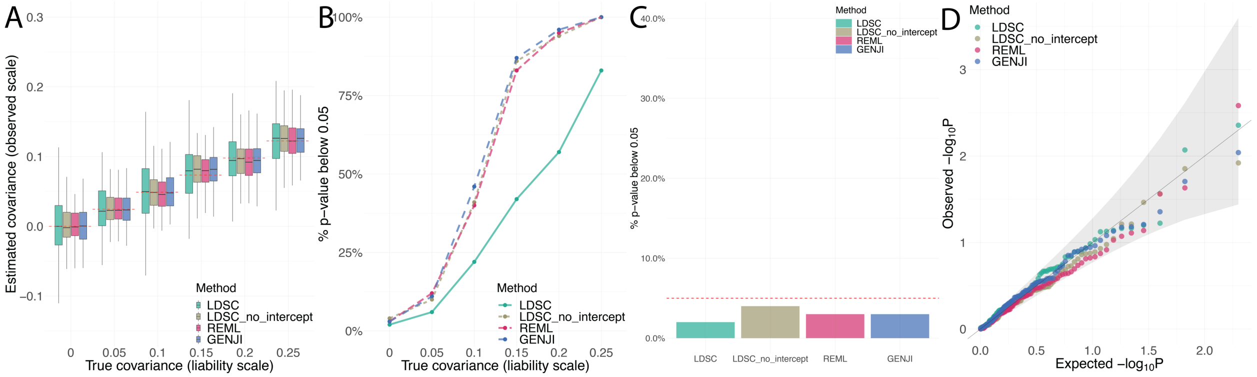

**Supplementary Figure 6. Evaluation of type I error control among transethnic genetic correlation (covariance) estimation methods for GWAS.** Popcorn using in-sample reference panel is denoted as "Popcorn_in_sample". The null hypothesis is corr=0 here. **(A)** Type I error when the true genetic correlation is 0. **(B)** qq-plot of p value when the true genetic correlation is set to be 0.

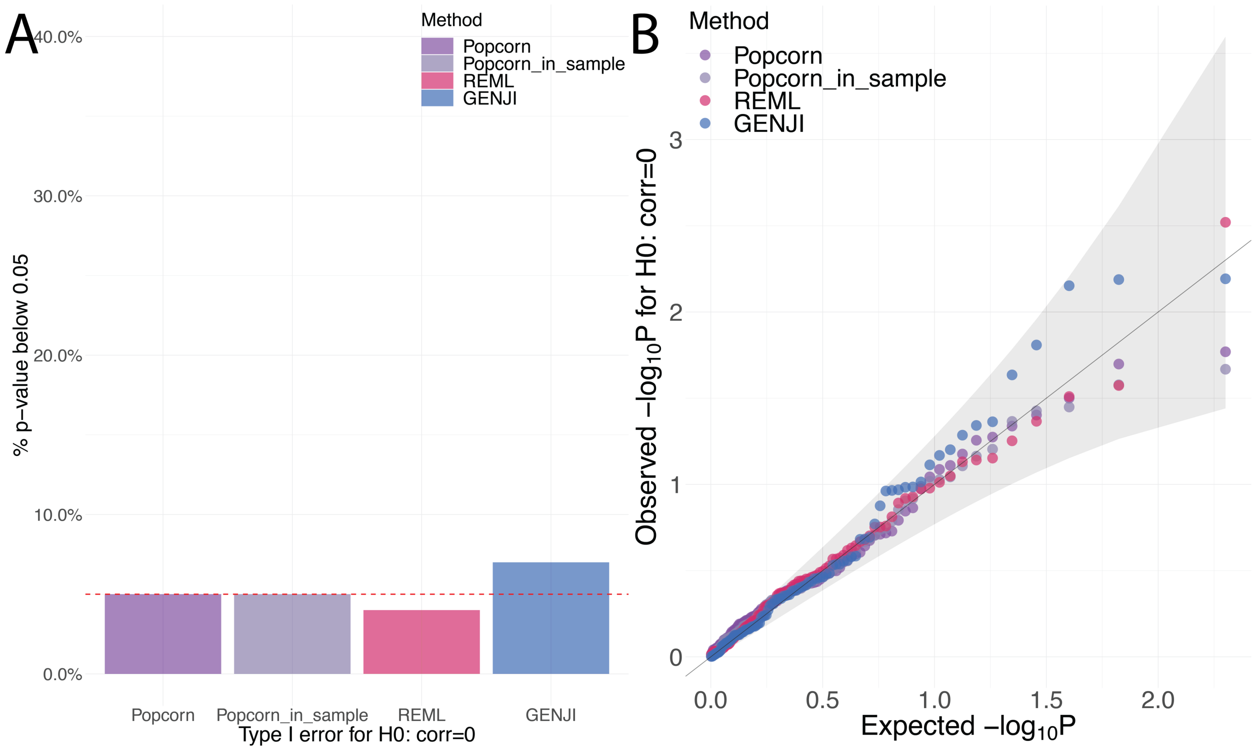

**Supplementary Figure 7. Evaluation of type I error control among transethnic genetic correlation (covariance) estimation methods for GWAS.** Popcorn using in-sample reference panel is denoted as "Popcorn_in_sample". The null hypothesis is corr=1 here. **(A)** Type I error when the true genetic correlation is 1. **(B)** qq-plot of p value when the true genetic correlation is set to be 1.

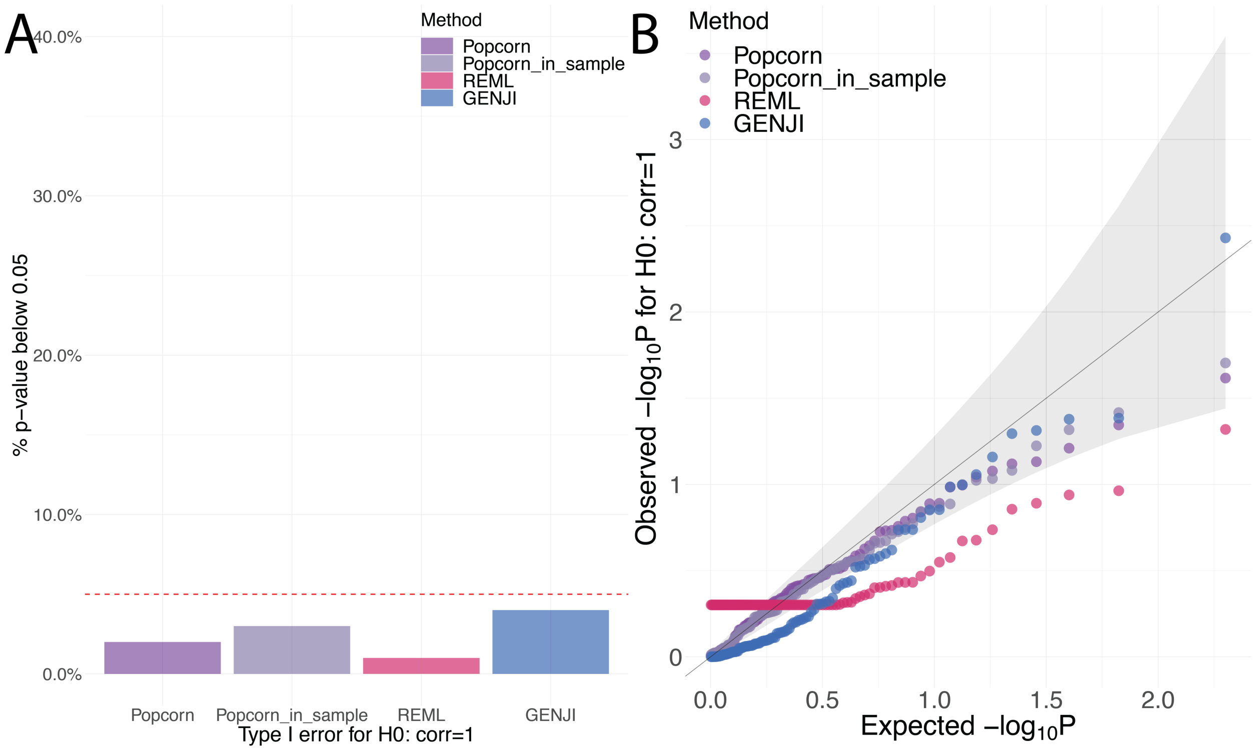

**Supplementary Figure 8. Comparisons of within-population genetic covariance and correlation estimation among REML, LDSC, and GENJI on WTCCC and NFBC traits.** The x-axis shows the estimates of REML while the y-axis shows the estimates of other methods. LDSC with intercept fixed at zero is denoted as "LDSC_no_intercept". The estimates of genetic **(A)** covariance and **(B)** correlation are summarized by scatter plots. The $R^2$ of LDSC, LDSC_no_intercept and GENJI for genetic covariance are 0.49, 0.92, and 0.92, respectively. The $R^2$ of LDSC, LDSC_no_intercept sand GENJI for genetic correlation are 0.43, 0.86, and 0.88, respectively. The color and shape of each point represent the method. The dashed lines are $y = x$.

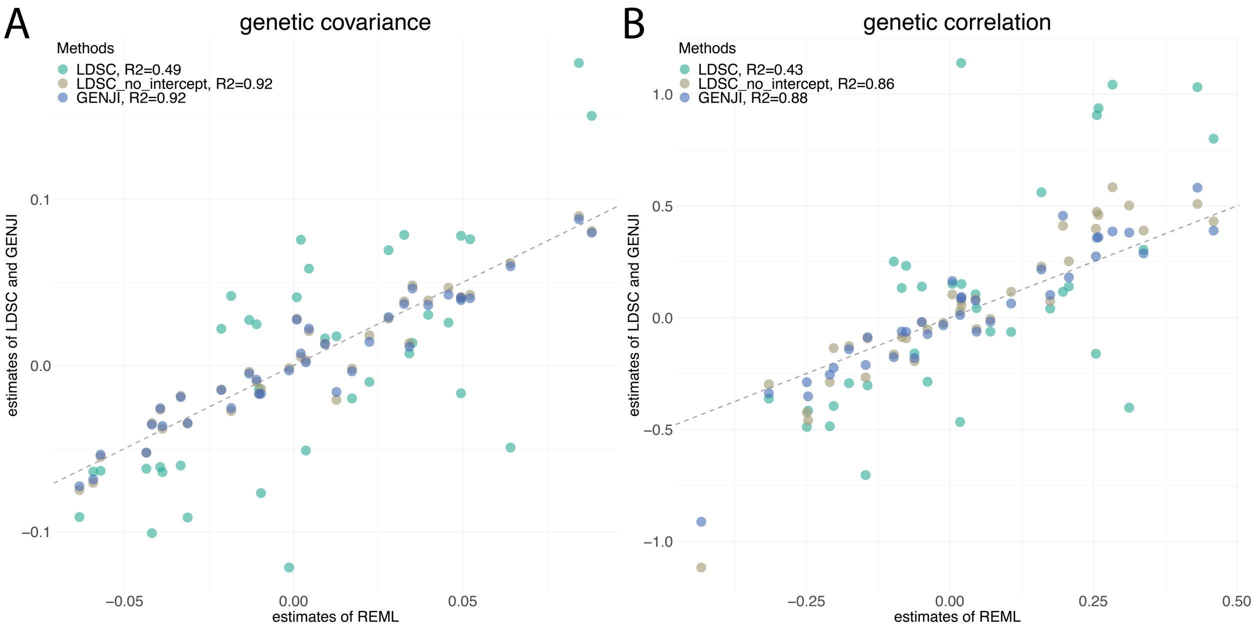

**Supplementary Figure 9. Comparisons of transethnic genetic covariance and correlation estimation among REML, Popcorn, and GENJI on UKBB traits of European and African populations.** The x-axis shows the estimates of REML while the y-axis shows the estimates of other methods. Popcorn using in-sample reference panel is denoted as "Popcorn_in_sample". The estimates of genetic **(A)** covariance and **(B)** correlation are summarized by scatter plots. The $R^2$ of Popcorn, Popcorn_in_sample and GENJI for genetic covariance are 0.93, 0.84, and 0.99, respectively. The $R^2$ of Popcorn, Popcorn_in_sample and GENJI for genetic correlation are 0.41, 0.45, and 0.94, respectively. The color and shape of each point represent the method. The dashed lines are $y = x$.

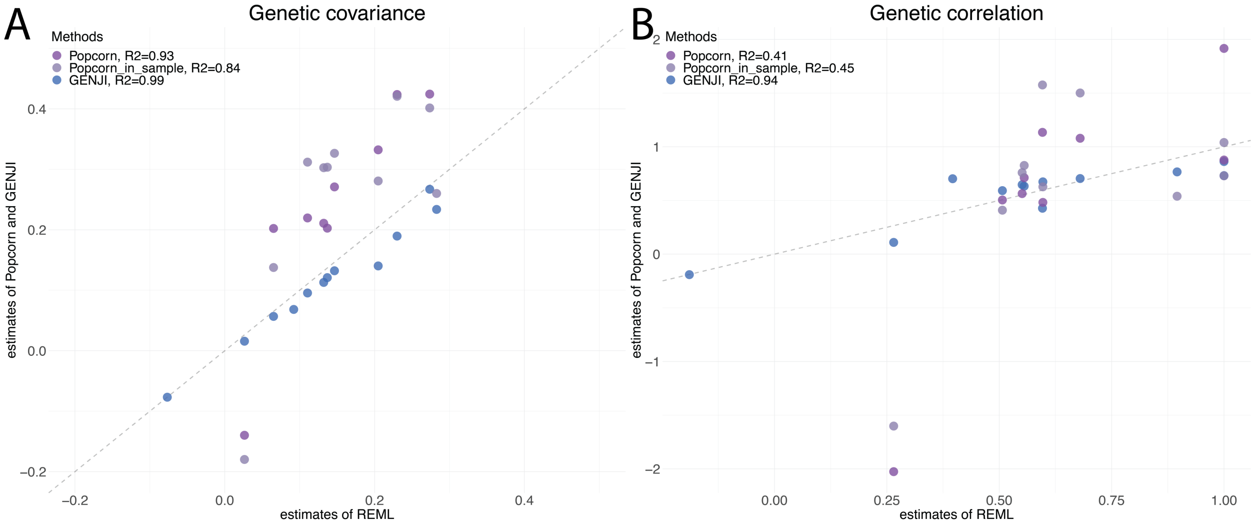

**Supplementary Figure 10. Transethnic genetic correlation estimates of REML, Popcorn and GENJI for 12 traits in UKBB between African and European populations.** To make the comparisons fair, we also include another implementation of Popcorn that use genotype data of African ancestry from UKBB as the reference panel of which GENJI takes the individual-level data as input. The point and the point range in the figures demonstrate the point estimates and their standard errors for transethnic genetic correlation of Popcorn and GENJI. Some of the estimates of Popcorn are not available due to low heritability estimation.

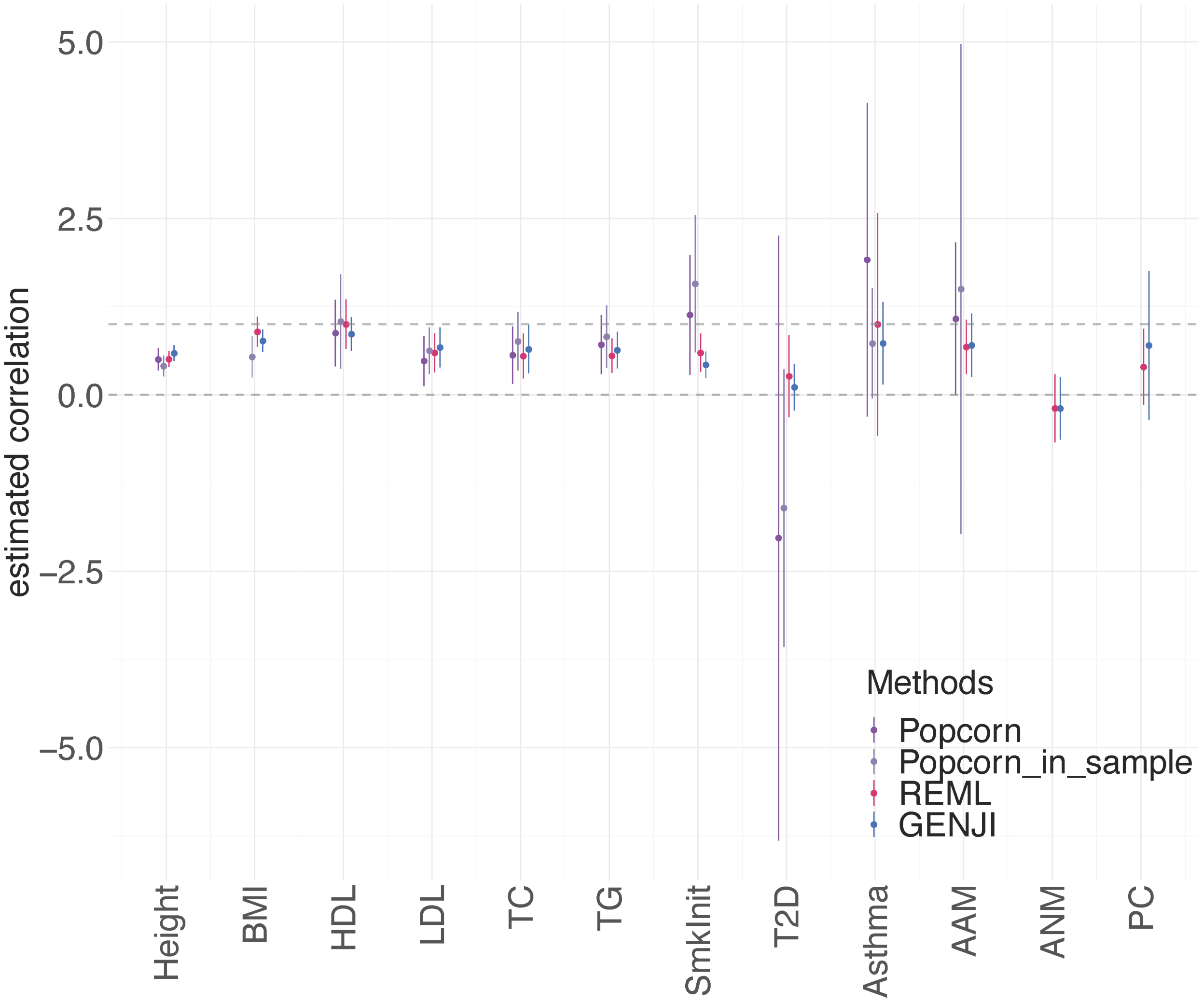
